# A Host-Harbored Metabolic Susceptibility of Coronavirus Enables Broad-Spectrum Targeting

**DOI:** 10.1101/2022.12.07.519404

**Authors:** Huan Fang, Yonglun Wang, Lu Liu, Kunlun Cheng, Pei Li, Ya Tan, Xingjie Hao, Miao Mei, Xinxuan Xu, Yuanhang Yao, Fuwen Zan, Linzhi Wu, Yuangang Zhu, Bolin Xu, Dong Huang, Chaolong Wang, Xu Tan, Zhaohui Qian, Xiao-Wei Chen

## Abstract

Host-based antivirals could offer broad-spectrum therapeutics and prophylactics against the constantly-mutating viruses including the currently-ravaging coronavirus, yet must target cellular vulnerabilities of viruses without grossly endangering the host. Here we show that the master lipid regulator SREBP1 couples the phospholipid scramblase TMEM41B to constitute a host “metabolism-to-manufacture” cascade that maximizes membrane supplies to support coronaviral genome replication, harboring biosynthetic enzymes including Lipin1 as druggable viral-specific-essential (VSE) host genes. Moreover, pharmacological inhibition of Lipin1, by a moonlight function of the widely-prescribed beta-blocker Propranolol, metabolically uncouples the SREBP1-TMEM41B cascade and consequently exhibits broad-spectrum antiviral effects against coronaviruses, Zika virus, and Dengue virus. The data implicate a metabolism-based antiviral strategy that is well tolerated by the host, and a potential broad-spectrum medication against current and future coronavirus diseases.

## Main text

The global pandemic of acute respiratory coronavirus disease (COVID-19), caused by SARS-CoV-2, has resulted in ∼600 million confirmed cases and >6.5 million deaths as of November 2022 (*1, 2*). One major lesson from the current pandemic centers on the constant mutation of the highly contagious coronavirus, which is projected to cohabit with the world population and may continue to evolve (*3*). Moreover, while antibodies or small molecule drugs directly targeting viral factors have been developed and approved for emergency uses, viral mutations may arise or even be selected to escape their inhibition (*4, 5*). Agents targeting the structural engagement between viral proteins and cell surface receptors, however, may also be outpaced by viral mutations. These counterplays by the viral pathogens therefore highlights the necessity of targeting host pathways, as a broad-spectrum antiviral strategy that bears much lower mutation rates and therefore less risks of drug resistance (*6, 7*). Despite such advantages, host-based antivirals must exploit and target vulnerabilities of viruses, *en fait* cellular processes that are essential to the virus but not to the host, to ensure both efficacy and safety.

Coronaviruses belong to a highly diverse family of enveloped positive strand RNA viruses that, upon entry into host cells, express and rapidly replicate their genomic RNA for self-amplification and new rounds of infection of the host (*6*). The currently-ravaging SARS-CoV-2 belongs to the beta-coronavirus (βCoV) subfamily, which also includes the well-studied model coronavirus murine hepatitis virus (*8*). A common requirement in the replication of RNA viruses, coronaviruses included, is the induction of the replication organelle (ROs), an elaborate membranous structure characterized by double membrane vesicles or spheres (DMV or DMS) (*6, 9-11*). These spatial platforms harbor and enrich specific viral replication factors and host proteins, and may protect the replicating RNA from degradation and surveillance by host innate immunity (*12*). Additionally, the membranous encapsulation might even shield viral factors from small molecular inhibitors. Hence, dissolving RO has been proposed as a host-based, broad-spectrum antiviral strategy, which may combine the effects of dissembling the viral replication machinery and increasing their exposure to disabling or degrading agents. Of note, ROs intricately connect with the host endoplasmic reticulum (ER), where the majority of membrane bilayers are being produced and assembled (*11, 13, 14*), thereby indicating the involvement of lipid metabolism in the host cells.

Several unbiased genetic screens have identified TMEM41B as a critical yet enigmatic host factor for multiple RNA viruses, including particularly SARS-CoV-2 (*15-18*). TMEM41B was recently identified as an ER-localized phospholipid scramblase, capable of catalyzing the trans-bilayer shuttle of bulk amphipathic phospholipids in an ATP-independent manner *in vitro* (*19-21*). Importantly, loss of TMEM41B *in vivo* leads to distorted ER membranes and failure in the ER retention of the master lipid regulator SREBP/SCAP, cumulating into severe tissue damages and metabolic dysfunction (*21*). Hence, while the exact role of TMEM41B in the viral life cycle remains to be fully established, the profound side effects caused by TMEM41B inactivation may nonetheless prevent direct targeting of the factor essential to both the virus and the host.

SREBPs are master transcription factors in lipid metabolism and promote the production of cholesterol, fatty acids and triglycerides (*22, 23*), and are extensively exploited by many viruses including the coronaviruses (*24-27*). The activation of SREBPs from their ER-resided precursors requires the transmembrane chaperone SCAP, in a manner tightly controlled by cholesterol or fatty acids to constitute a delicate negative feedback loop (*23*). Of note, SREBP in Drosophila is negatively regulated by phospholipids, thereby implicating SREBP/SCAP as a transmembrane sensor for ER membrane structure and integrity (*28, 29*). Nevertheless, the role of SREBPs in phospholipid metabolism and especially membrane biogenesis is much less understood, compared to those of cholesterol or fatty acids (*30, 31*).

Here we show that SREBP1 and TMEM41B constitute a host cascade to couple phospholipid synthesis (metabolism) and bilayer assembly (manufacture), thereby maximizing membrane supplies to house viral replication. Serendipitously, the metabolism-to-manufacture cascade also exposes a metabolic vulnerability of coronaviruses, harboring enzymes such as Lipin1 that is required for the life cycle of viruses but not of host cells. Consequently, the widely-prescribed medicine Propranolol can be employed to inhibit Lipin1 and uncouple the SREBP1-TMEM41B cascade, producing broad spectrum antiviral effects via disrupting RO formation. Propranolol effectively counters multi-tissue pathogenesis in murine models of coronavirus infection, implicating an accessible and broad-spectrum treatment option against coronaviruses at the current or future times.

## Results

### Direct regulation on TMEM41B by SREBP1 constitutes a membragenic cascade

We first sought to delineate the regulations on the recently identified phospholipid scramblases, which could equilibrate the intrinsic asymmetry resulted from the leaflet-specific synthesis of phospholipids (*32, 33*). Interestingly, immunoblotting (IB) showed that active SREBP1 selectively upregulated TMEM41B, but not related enzymes such as TMEM41A or VMP1 (Fig. 1A, quantified in 1B). The upregulation of TMEM41B was comparable to established SREBP1 targets in lipid biosynthesis, including fatty acid synthase (FASN) and the phosphatic acid phosphatase Lipin1, implicating the scramblase as a *bona fide* target of the master transcription factor of lipid metabolism. Accordingly, active SREBP1 selectively upregulated mRNA levels of TMEM41B, but not TMEM41A or VMP1, also at levels comparable to FASN (Fig. 1C). Collectively, the data suggested an unexpected regulation from the very upstream regulator SREBP1 to a specific scramblase catalyzing the final step in bilayer manufacture.

**Figure 1.**
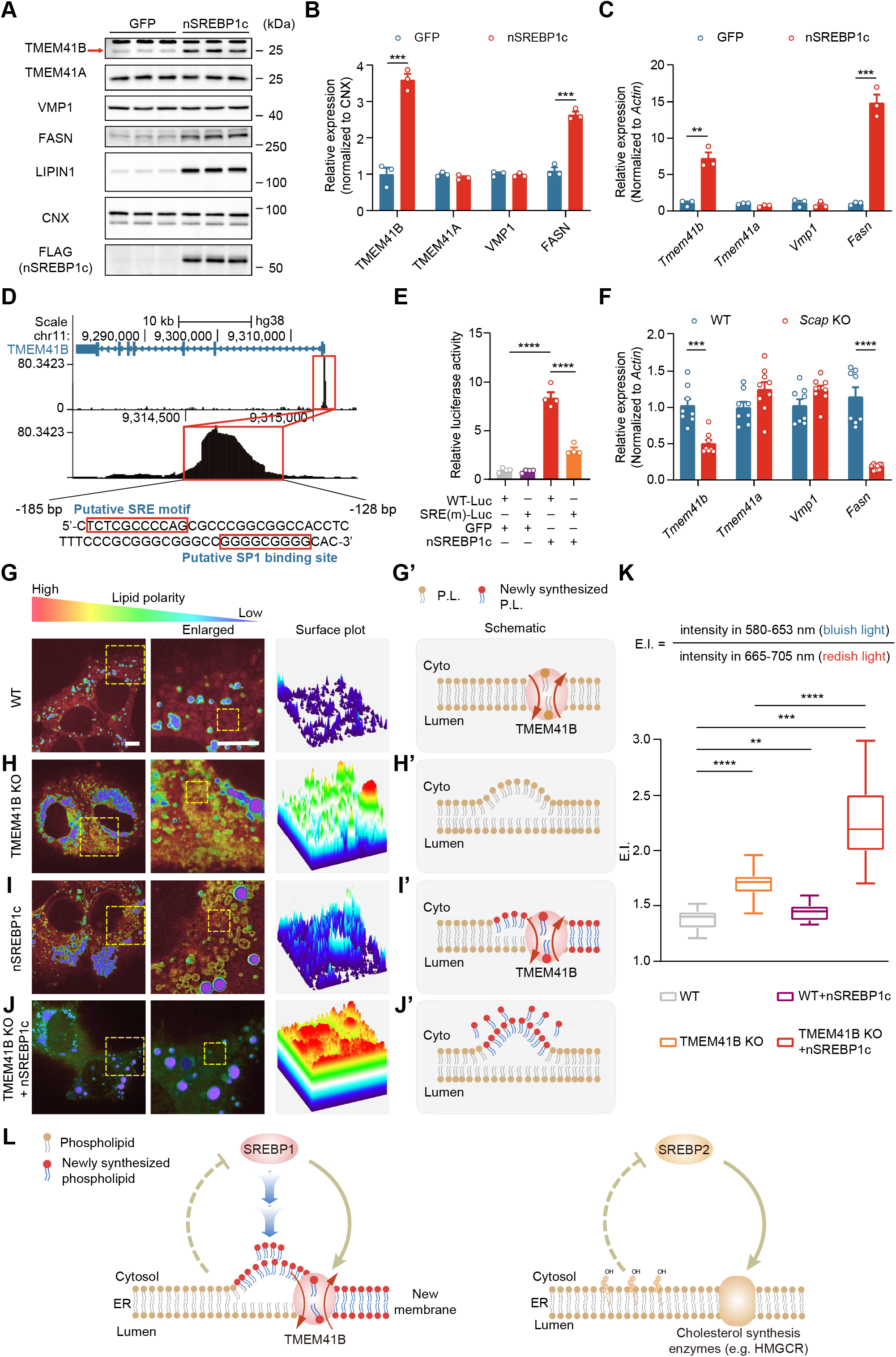
Direct regulation on TMEM41B by SREBP1 constitutes a membragenic circuit. (A) Active SREBP1 upregulates TMEM41B protein. Primary hepatocytes infected with AAV expressing GFP (control) or active SREBP1 are subjected to cell lysis and immuno-blotting (IB) with antibodies against the indicated proteins. Representative IB from 4 independent experiments are shown. (B) Quantification of IB in (A). Data are presented as mean ± SEM. ***, p < 0.001, ****, p < 0.0001. (C) Active SREBP1 upregulates *Tmem41b* transcripts. Primary hepatocytes infected with AAV expressing GFP (control) or active SREBP1 are subjected to RNA harvest and qPCR for the indicated genes. Representative experiments from 4 independent experiments are shown. Data are presented as mean ± SEM. **, p < 0.01, ***, p < 0.001. (D) SREBP1 occupancy on the promoter of *TMEM41B* gene revealed by chromatin-immunoprecipitation and sequencing (CHIP-Seq) datasets. Upper and middle: genome browser view of the *TMEM41B* gene and SREBP1 CHIP-seq peaks. Lower: enlarged view of the SREBP1-occupied regions. SREBP1 binding motif (left) and SP1 binding motif are highlighted by the red rectangles, respectively. (E) Human active SREBP1 binds to the SRE in *TMEM41B* promoter. *TMEM41B* promoter or SRE mutant promoter are cloned in the luciferase reporter vector, and transfected into 293A cells together with Renilla, with active SREBP1 or GFP. Data are presented as mean ± SEM. ****, p < 0.0001. (F) Loss of hepatic *Scap* down-regulates *Tmem41b* transcripts. qPCR results showing transcription levels of *Tmem41b* in *Scap*^*flox/flox*^ mice receiving AAV-TBG-GFP (n = 8) or AAV-TBG-CRE (n = 8). Representative experiments from 3 independent experiments are shown. Data are presented as mean ± SEM. ***, p < 0.001, ****, p < 0.0001. (G) Ratiometric images and the surface plot depicting the emission index of wild type Huh7 cells. Cells are labelled with Nile Red and subjected to live cell super resolution microscopy, collecting emission signals from the red and the blue channels (n = 3 independent experiments). Right: the ratiometric spectrum of the enlarged field. Scale bar=5 μm. Model depicted in (G’). (H) Ratiometric images and the surface plot of TMEM41B-deficient Huh7 cells. Model depicted in (H’). (I) Ratiometric images and the surface plot of wild type Huh7 cells receiving active SREBP1. Model depicted in (I’). (J) Ratiometric images and the surface plot of TMEM41B-deficient Huh7 cells receiving active SREBP1. Model depicted in (J’). (K) Quantification of mean emission indexes in cells from (G-J). Data are presented as mean ± SEM. **, p < 0.01, ****, p < 0.0001. (L) Schematic diagram of the SREBP1-TMEM41B circuit that couples phospholipid production and bilayer assembly to drive membragenesis (left), and the established SREBP2 circuit in cholesterol metabolism and homeostasis (right).

To delineate the molecular basis by which SREBP1 regulates TMEM41B, we unbiasedly surveyed public chromatin-immunoprecipitation and sequencing (CHIP-Seq) datasets. The data revealed SREBP1 occupancy (*in-trans*) at the promoter regions of the *TMEM41B* gene, which contain DNA sequences (*in-cis*) resembling the sterol-responsive-element (SRE) consensus of SREBP1 (Fig. 1D). Moreover, active SREBP1 upregulated the expression of the luciferase reporter fused with wild type *TMEM41B* promoter, whereas the upregulation became blunted when the SRE motif was mutated (Fig. 1E). To further confirm the role of SREBP in TMEM41B regulation, we employed mice lacking hepatic SCAP, the transmembrane chaperone required for the ER-Golgi escorting and therefore activation of SREBP. Consistent with the gain-of-function data, mRNA levels of *Tmem41b* or *Fasn*, but not *Tmem41a* or *Vmp1*, were diminished in *Scap*^*-/-*^ livers compared to wild type controls (Fig. 1F).

The direct regulation of TMEM41B by SREBP1 suggest that, in response to demands for elevated membrane biogenesis, the intrinsic imbalance between bilayer leaflets needs to be resolved by adequate lipid scrambling. To test this hypothesis, we took advantage of the advances in ratiometric spectrum imaging that combined live cell super resolution microscopy with the biocompatible and solvatochromic nature of the lipid dye Nile Red (*34, 35*). The dye labels intracellular membranes and lipid storages, exhibiting blue shifts in its emission spectrum in a less polar environment in membrane organization (Fig. S1). As reported previously (*34*), ratiometric depiction of the emission indexes (EI) from the red and the blue channels confirmed that the ER membranes and the core of the lipid droplets represented the lowest and highest values in wild type Huh7 cells (Fig. 1G, model depicted in G’), respectively. By contrast, TMEM41B-deficent cells displayed blue-shifted emission index in the ER regions compared to wild type controls (Fig. 1H, depicted in H’), suggesting that defective lipid scrambling caused unmatched bilayer leaflets with decreased polarity of local lipid environment.

Interestingly, induction of lipid biogenesis by ectopic expression of active SREBP1 caused modest yet significant blue shifts in the emission index of the ER (Fig. 1I, quantified in 1K), further suggesting that leaflet imbalance is a general phenomenon during membrane biogenesis and needs to be adequately equilibrated by lipid scrambling. Consistent with this notion, SREBP1 expression synergized with TMEM41B deficiency to cause large blue shifts in the emission (Fig. 1J, quantified in 1K), revealing greater unmatching of ER bilayers caused by an elevation of asymmetric phospholipid flux plus defective lipid scrambling (also depicted in 1J’).

Taken together with previous reports that loss of TMEM41B causes drastic ER membrane distortion and uncontrolled SREBP activation (*21*), the results revealed a feedback circuit between SREBP1 and TMEM41B in phospholipid metabolism (Fig. 1L, left). Moreover, this novel regulatory scheme appears to mirror the well-established SREBP-dependent regulatory circuit that robustly controls cholesterol metabolism (*22, 23*) (also depicted in Fig. 1L, right). Therefore, by coupling phospholipid *production* and bilayer *assembly*, the SREBP1-TMEM41B cascade could maximize membrane supply, together streamlining a “metabolism-to-manufacture” process.

### Host membrane manufacture exposes a metabolic susceptibility of coronaviruses

The functional pairing of two essential host factors, SREBP1 and TMEM41B, pointed to a mechanism that could be exploited by the virus for maximizing membrane supply from the host, through the hijacking of host phospholipid production and the subsequent bilayer assembly (Fig. 2A). Consistent with this “metabolism-to-manufacture” hypothesis with respect to the host SREBP1-TMEM41B cascade in viral life cycle, ratiometric spectrum imaging revealed substantial membrane alterations in primary hepatocytes infected with the model beta-coronavirus (βCoV) murine hepatitis virus (Fig. 2B, quantified in 2C). The data thus implied an upregulation of phospholipid production in need of bilayer equilibration by coronaviruses. Consistent with this notion, ectopic supply of phospholipids by ER-targeted liposomes or expression of active SREBP1 both led to an increase in βCoV loads in wild type cells compared to controls, suggesting that phospholipid supply represents a limiting factor for coronavirus amplification in cells (Fig. 2D, left three columns). In cells lacking TMEM41B, however, viral production became absent in control, phospholipid-supplied, or SREBP1-expressed conditions (Fig. 2D, right three columns). These data therefore confirmed the requirement of bilayer assembly downstream of the metabolic supply of phospholipids to support viral life cycle, together orchestrated by the host SREBP1-TMEM41B cascade.

**Figure 2.**
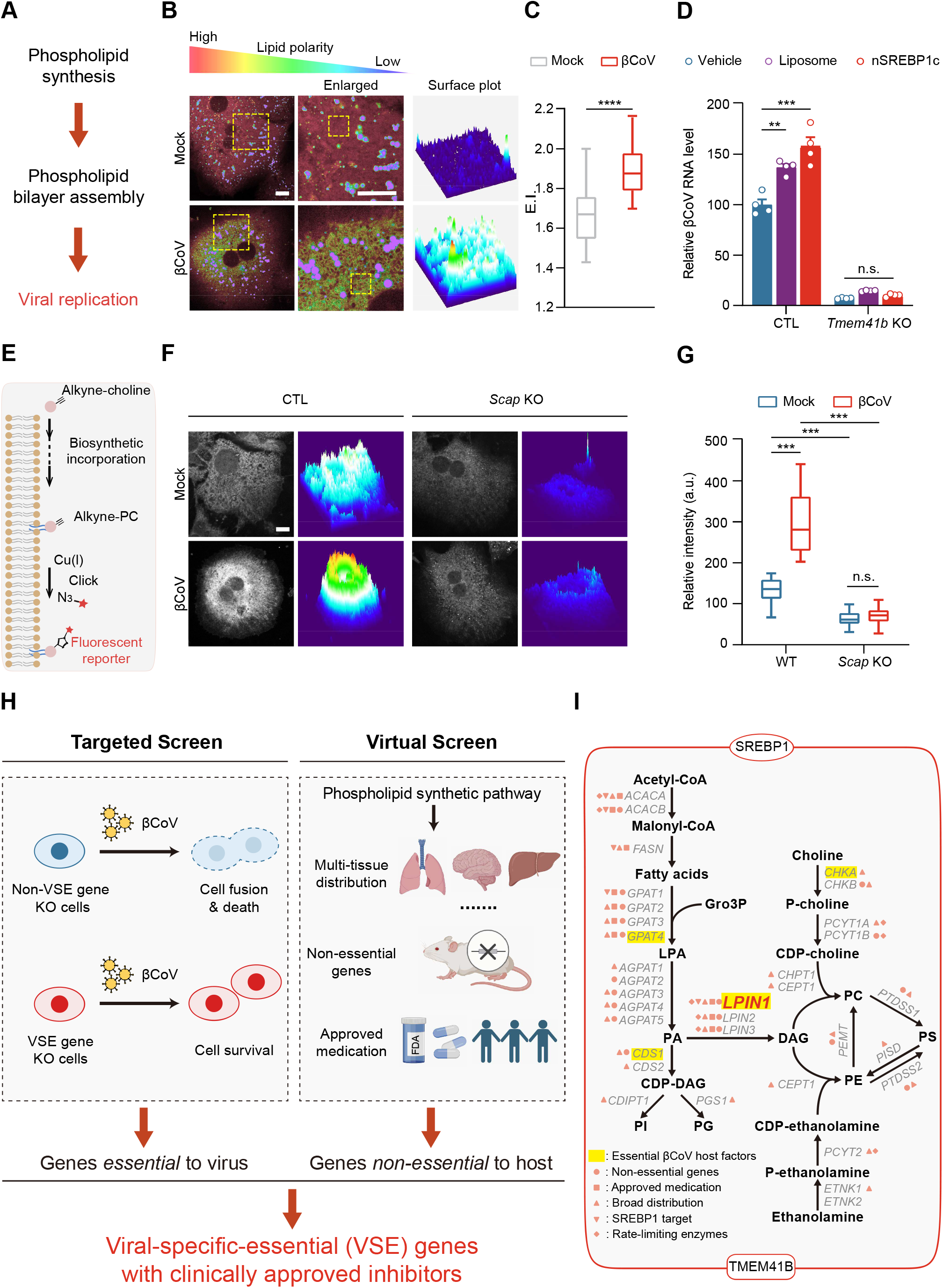
Host membragenic circuit exposes a metabolic susceptibility of coronaviruses. (A) Schematic diagram of the “membragenic hypothesis” by which host SREBP1-TMEM41B supplies phospholipids and membrane bilayer for viral replication. (B) Ratiometric images of βCoV-infected primary hepatocytes showing substantial spectrum shifts compared to uninfected control. Representative results from 3 independent experiments are shown. Scale bar = 5 μm. (C) Quantification of mean emission indexes from (B). Data are presented as mean ± SEM. ****, p < 0.0001. (D) Phospholipid supply or ectopic active SREBP1 expression promotes βCoV production, and their blockade by *Tmem41b* KO. Wild type (blue columns) or *Tmem41b* KO (red columns) hepatocytes receiving the indicated agents are infected, prior to RNA harvest and qPCR for viral N transcripts. Representative qPCR results from 3 independent experiments are shown. Data are presented as mean ± SEM. n.s., no significance. **, p < 0.01, ***, p < 0.001. (E) A chemical-biology approach to trace phospholipid production in cells. Cells are metabolically labeled with alkyne-choline to allow biosynthetic incorporation into alkyne-PC, which can be conjugated with fluorophores via click-chemistry for microscopic visualization. (F) βCoV infection activates phospholipid production in cells. Primary hepatocytes are left uninfected or infected with βCoV, labeled as in (E) prior to visualization. Left: confocal images. Right: surface plots of alkyne-PC signals. Scale bar = 5 μm. Representative results from 3 independent experiments are shown. Scale bar = 5 μm. (G) Quantification of alkyne-PC signals from control or βCoV infected wild type or *Scap* KO hepatocytes. Data are presented as mean ± SEM. n.s., no significance. ****, p < 0.0001. (H) The scheme of the hybrid screen which identifies <u>v</u>iral-<u>s</u>pecific <u>e</u>ssential (VSE) genes. (I) Summary of the screen results in (H) and the identification of *Lpin1* as a <u>v</u>iral-<u>s</u>pecific <u>e</u>ssential (VSE) gene. Essential host factors are highlighted by yellow, whereas genes are remarked by virtue of having clinically approved inhibitors (square), broad distributed (triangle), SREBP1 targets (inverted triangle), or rate-limiting enzymes (rhombus), respectively.

The above data further indicated a metabolic flux of phospholipids towards the TMEM41B scramblase that generates bilayer membranes to support viral life cycle. To test this hypothesis, we adapted a “click-chemistry” based chemo-metabolic labelling approach to track the production of phosphatidylcholine (PC), the most abundant form of membragenic phospholipids in mammalian cells (*33*). Figure 2E illustrates that choline with a two-carbon alkyne tag (Alkyne-choline) can be converted into PC at the ER through the Kennedy pathway, and the newly-synthesized PC may be visualized after biorthogonal conjugation with fluorophores in cells. Of note, βCoV infection caused a significant upregulation of alkyne-PC signal compared to un-infected cells, reflecting an induction of phospholipid production that may support viral replication (Fig. 2F, quantified in 2G). Alkyne-PC signals became reduced in *Scap*-deficient hepatocytes, supporting the role of SREBP in phospholipid production. Importantly, βCoV-induced alkyne-PC signals were blunted upon loss of SCAP, further implicating SREBPs in the induction of membragenesis triggered by coronaviruses (Fig. 2F&G).

The upregulation of host phospholipid metabolism by coronaviruses culminated into the possibility of limiting the metabolic flux through the host SREBP1-TMEM41B cascade to counter the viral pathogen, furthering that phospholipid enzymes within the cascade may be required for the life cycle of viruses but not of the host. To identify such <u>v</u>iral-<u>s</u>pecific <u>e</u>ssential (VSE) genes harbored within the host SREBP1-TMEM41B cascade, we designed a targeted CRISPR screen based on the cell lethality caused by coronavirus infection. In essence, we hypothesize that loss of VSE genes could generate grossly healthy “mutant” cells that could escape the coronavirus-caused fusion and eventually death in control cells (Fig. 2H, left). Moreover, as coronavirus often induces cell syncytia and likely collateral damage to healthy host cells (Fig. S2), we reason that a targeted screen would generate a homogeneous cell population that allow direct inspection of cell health, while avoiding the loss of the coronavirus-resistant cells as an under-represented population in pooled screens due to potential collateral damages from the coronavirus-sensitive cells. In parallel to the targeted genetic screen in cells, we also performed a virtual screen to search for host factors that are essential for the virus, but not to the host cells (Fig. 2H, right). Such “ideal” host factor, however, would need to be broadly distributed to counter the multiple tissue pathogenesis of coronaviruses and have clinically approved targeting agents that support the safety while allowing subsequent application.

Interestingly, the targeted CRISPR screen identified that the SREBP1-TMEM41B cascade was enriched with biosynthetic enzymes as essential host factors for βCoV (Fig. 2I, highlighted by yellow), implicating the membrane manufacture cascade as a metabolic susceptibility of the virus. Loss of these metabolic genes (including GPAT4, CDS1, LIPIN1, and CHKA) abolished βCoV-induced syncytia formation and death in human lung A549 cells stably expressing the βCoV receptor (Fig. S2B). When combined with the virtual screen, LIPIN1 stood out as a druggable VSE gene among these newly-identified host factors, as a broadly expressed yet non-essential gene in either human or mice (Fig. S3). The pharmacological targeting of the Lipin1 enzyme by clinically approved medicines with respect to viral inhibition will be investigated in the later part of the study. Collectively, the data suggested that the SREBP1-TMEM41B cascade drives membrane production to constitute a metabolic susceptibility of coronavirus, and importantly, the biosynthetic enzyme Lipin1 within the cascade may represent a VSE factor that can be safely targeted to counter the viruses without endangering the host.

### Lipin1 inhibition targets metabolic susceptibility of coronaviruses

Consistent with the above screening experiments in A549 cells, loss of Lipin1 but not Lipin2 also diminished βCoV load in primary hepatocytes, as did with *Scap* or *Tmem41b* ablation (Fig. 3A). Unlike SCAP or TMEM41B, however, Lipin1 inactivation didn’t appear to decrease cell viability or fitness over time (data not shown), consistent with previous reports that even complete deficiency of the gene didn’t cause lethality from yeast to humans (*36, 37*). These results together reflected the non-essential nature of the enzyme in the host, thus allowing its inhibition for antiviral applications safely. Biochemically, Lipins dephosphorylate phosphatic acid (PA) to diacylglycerol (DAG) as precursors for the production of PC and PE, together accounting for ∼80-90% of membragenic phospholipids in cells (*37, 38*), further suggesting a unique dependence of membrane supply in the viral life cycle. Consistent with this notion, while Lipin1 deficiency diminished βCoV loads in primary hepatocytes, phospholipid replenishment via liposomes largely rescued the inhibition (Fig. 3B). Taken together, the data suggested that phospholipid production and membrane manufacture operated by the SREBP1-TMEM41B cascade represents a limiting resource for the life cycle of coronaviruses, and importantly, Lipin1 ablation could metabolically block the host process to counter the viral pathogen.

**Figure 3.**
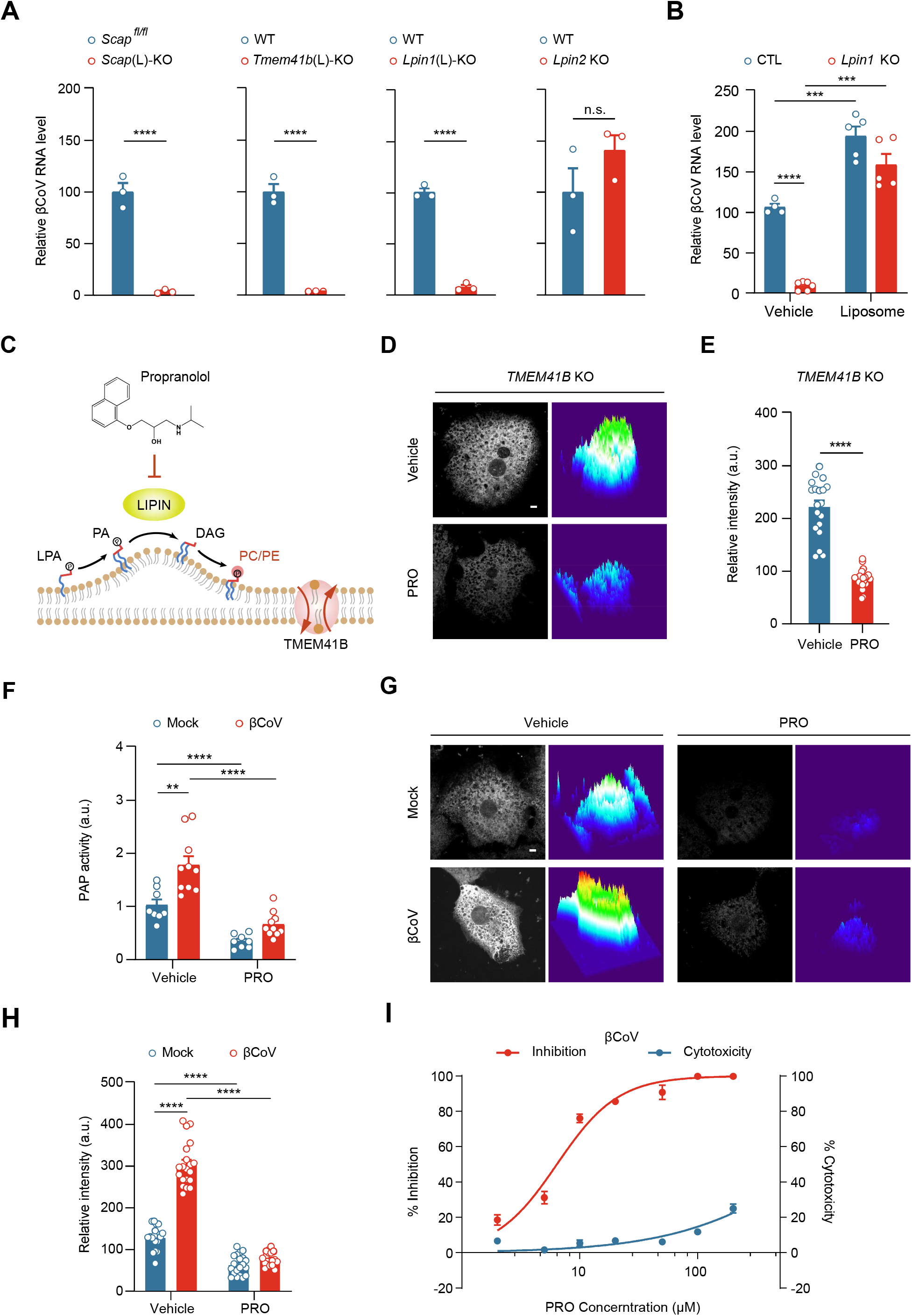
Lipin1 inhibition by Propranolol enables a broad spectrum antiviral strategy. (A) Loss of *Scap, Tmem41b* or *Lpin1* diminishes βCoV load in cells. Primary hepatocytes isolated from mice with the indicated genotypes are infected with βCoV, prior to RNA harvest and qPCR for viral N transcripts. Representative results from 3 independent experiments are shown. Data are presented as mean ± SEM. ****, p < 0.0001. (B) *Lpin1* ablation blocks the production of βCoV and its rescue by phospholipid addition. Control or hepatic *Lpin1* KO hepatocytes are infected with βCoV, and added 60 μM liposomes at 2 h.p.i, prior to RNA harvest for RT-qPCR. Representative results from 3 independent experiments are shown. Data are presented as mean ± SEM. ***, p < 0.001, ****, p < 0.0001. (C) Schematic diagram of Kennedy pathway for phospholipid (PC/PE) production at the ER, centered on the catalysis by LIPIN that dephosphorylates phosphatidic acid (PA) to produce diacylglycerol (DAG) and its inhibition by Propranolol. (D) Propranolol (PRO) decreases alkyne-PC accumulation in TMEM41B KO Huh7 cells. Cells receiving vehicle or 50 μM Propranolol are labeled by alkyne-choline prior to click conjugation and confocal microscopy. Representative results from 3 independent experiments are shown. Left: confocal images. Right: surface plots. Scale bar=5 μm. (E) Quantification of alkyne-PC signals from (D). Data are presented as mean ± SEM. ****, p < 0.0001. (F) βCoV infection activates cellular PAP enzyme activity that can be inhibited by Propranolol. Primary hepatocytes receiving vehicle or 50 μM Propranolol are uninfected or infected with βCoV. PA and DAG levels are quantified in lipid extracts by LC-MS. Data are presented as mean ± SEM. **, p < 0.01, ****, p < 0.0001. A.U., arbitrary unit. (G) Propranolol decreases alkyne-PC production induced by βCoV infection. Primary hepatocytes receiving vehicle or 50 μM Propranolol are uninfected or infected with βCoV. Cells are labeled by alkyne-choline prior to click conjugation and confocal microscopy. Representative results from 3 independent experiments are shown. Left: confocal images. Right: surface plots. Scale bar=5 μm. (H) Quantification of alkyne-PC signals from (G). Data are presented as mean ± SEM. ****, p < 0.0001. (I) Dosage-independent anti-βCoV effects by Propranolol. Primary hepatocytes treated for 12 h with indicated doses of Propranolol are infected with βCoV, prior to RNA harvest for RT-qPCR. Drug cytotoxicity is measured via CCK8 assay. Representative results from 3 independent experiments are shown. Data are presented as mean ± SEM.

Of particular note, the phosphatic acid phosphatase (PAP) activity of Lipin can be effectively inhibited by the lipophilic beta-blocker Propranolol as a moonlight function (Fig. 3C) (*39-42*). Indeed, Propranolol diminished the accumulation of Alkyne-choline signal upon TMEM41B deficiency (Fig. 3D, quantified in 3E), further confirming that the drug uncoupled the SREBP1-TMEM41B cascade by limiting the metabolic flux of membragenic phospholipids. Propranolol has been widely used to treat cardiovascular complications with minimal toxicity, and has also been repurposed for a number of novel therapeutic indications (*43-46*). These applications further suggest that inhibition of Lipin could be well tolerated in humans in clinically relevant manners, leading us to examine the anti-coronaviral effects with Propranolol in cells. Consistent with an active requirement of membragenesis in coronavirus infection, βCoV induced an increase in cellular PAP activity, which was blunted by Propranolol (Fig. 3F). As seen with genetic ablation of *Scap*, metabolic tracing following Alkyne-choline conversion to Alkyne-PC also revealed that Propranolol diminished the production of PC in primary hepatocytes induced upon βCoV infection (Fig. 3G, quantified in 3H). Consistent with the required role of SREBP1-TMEM41B and membragenesis for viral life cycle, Propranolol decreased βCoV load in primary hepatocytes in a dose-dependent manner, with IC_50_ at 6.39 μM (Fig. 3I, red line). By contrast, Propranolol exhibited little effects on βCoV binding to cell surface or entry into hepatocytes (data not shown), nor did it cause overt cytotoxicity up to 100 μM (Fig. 3I, blue line). Taken together, the results suggested that Propranolol effectively exploited the viral vulnerability harbored in the host SREBP1-TMEM41B cascade, thereby countering coronavirus by limiting the metabolic flux and blocking membrane manufacture that are required by the viral pathogen.

### Metabolic blockade by Propranolol enables a broad-spectrum antiviral strategy

The above antiviral effects intrigued us to directly examine the impact on the viral life cycle upon Propranolol-enabled metabolic blockade, by performing volumetric tomography on coronavirus-infected cells with the focus ion beam coupled scanning electron microscopy (FIB-SEM). As previously established (*9-11*), βCoV infection in primary hepatocytes potently triggered the formation of the membranous replication compartments, featured with the characteristic double membrane enclosed vesicles (DMV) and double membrane enclosed smaller spheres (DMS) (Fig. 4A, left and enlarged panels, and movie S1). These ROs were often in conjunction with the host ER membranes which have also undergone drastic morphological changes. Strikingly, Propranolol treatment largely eliminated DMV or DMS structures induced by coronavirus infection (Fig. 4A, right panel, and movie S2), and maintained the reticular ER structures as also visualized by thin-sectioning and transmission EM (Fig. S4). Taken together, the tomographic and EM analysis revealed an unexpected sensitivity of RO formation to the metabolic blockade by Propranolol, consistent with the notion of a high demand of membrane supply in constructing these viral replication compartments in cells.

**Figure 4.**
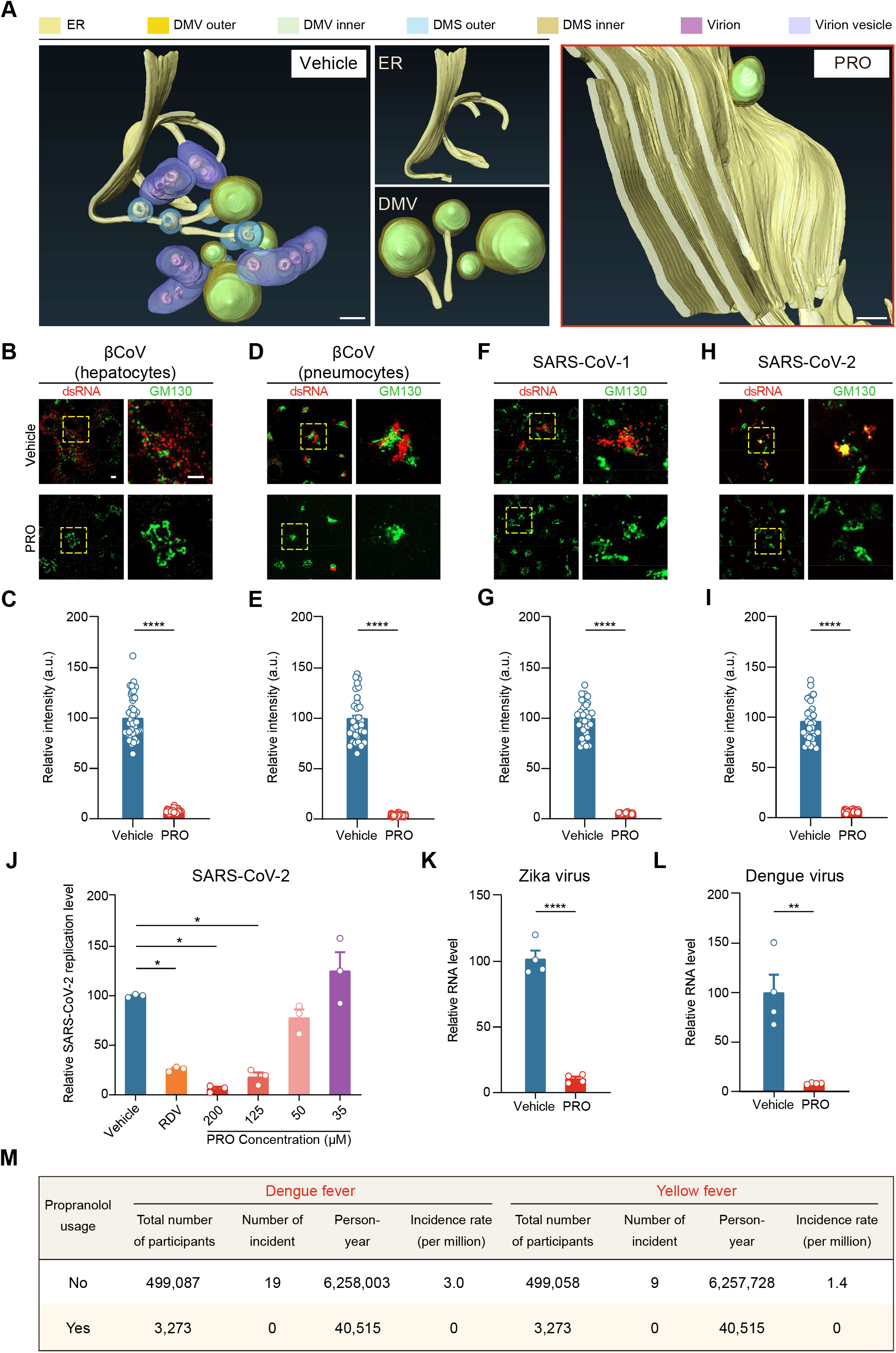
Metabolic blockade by Propranolol enables a broad-spectrum antiviral strategy. (A) Volumetric tomography of replication organelles (ROs) induced by βCoV infection and their elimination by Propranolol. Hepatocytes infected with βCoV are subjected to FIB-SEM and tomographic reconstruction. Upper: coronavirus induces ROs formation. Lower: lack of viral ROs in βCoV infected cells receiving 50 μM Propranolol. Scale bar=200 nm. (B) Propranolol exhibits broad spectrum anti-coronavirus ability in cells. Primary hepatocytes pretreated with vehicle or 50 μM Propranolol for 12 h are infected by MHV for 8 h, prior to immunofluorescence. Scale bar=5 μm. (C) Quantification of (B), ****, p < 0.0001. (D) Confocal images of dsRNA in A549-mCEACAM1 cells pretreated with vehicle or 50 μM Propranolol for 12 h prior to MHV infection for 8 h. (E) Quantification of (D), ****, p < 0.0001. (F) Confocal images of dsRNA in HEK293T cells pretreated with vehicle or 125 μM propranolol for 6 h prior to being transfected with SARS-CoV-1 replicon for 24 h. (G) Quantification of (F), ****, p < 0.0001. (H) Confocal images of dsRNA in HEK293T cells pretreated with vehicle or 125 μM propranolol for 6 h prior to being transfected with SARS-CoV-2 replicon for 24 h. (I) Quantification of (H), ****, p < 0.0001. (J) Propranolol blocks SARS-CoV-2 replication in cells. HEK293T cells expressing SARS-CoV-2 replicons are treated with indicated doses (μM) of Propranolol or Remdesivir (as the positive control), prior to cell harvest and luciferase assay. Representative results from 3 independent experiments are shown. Data are presented as mean ± SEM. *, p < 0.05. (K) Propranolol inhibits ZIKV production in U87MG cells. U87MG cells are pre-treated with 20 μM Propranolol for 12h, infected with MOI = 0.1 ZIKV, harvested for RT-qPCR at 48 h post infection. Representative results from 3 independent experiments are shown. Data are presented as mean ± SEM. ****, p < 0.0001. (L) Cellular DENV RNA is determined as in (K). Data are presented as mean ± SEM. **, p < 0.01. (M) Real world data from the UK Biobank on Propranolol usage and incidence of Dengue fever and Yellow fever.

ROs represent a common requirement for RNA replication of coronaviruses and flaviviruses in different cells, thus leading us to hypothesize that Propranolol may exhibit broad-spectrum antiviral effects. Indeed, Propranolol treatment diminished dsRNA of βCoV in both primary hepatocytes (Fig. 4B, quantified in 4C) and A549 pneumocytes (Fig. 4D&E). Moreover, viral dsRNA generated by the replicons of SARS-CoV-1 (Fig. 4F&G) and SARS-CoV-2 (Fig. 4H&I) were also eliminated by Propranolol treatment, as with luciferase reporter of SARS-CoV-2 in HEK293 cells (Fig. 4J). Lastly, Propranolol also depleted virus load of Zika virus (Figure 4K) and Dengue virus (Fig. 4L) after infection of U87MG cells, respectively. Interestingly, a survey of the real world data from the UK Biobank also revealed diminished incidents of Dengue fever or yellow fever in people taking Propranolol, though infections of these flaviviruses were rather rare in general (Fig. 4M). Collectively, the biochemical, ultra-structural, and functional data demonstrated the broad-spectrum antiviral effects by the host-based drug, further suggesting the feasibility of targeting the membragenic process driven by the SREBP1-TMEM41B cascade for viral defense.

### Propranolol exhibits broad protection against coronavirus pathology in mice

The above cellular data led us to examine the antiviral effects of Propranolol-enabled metabolic blockade of the host SREBP1-TMEM41B cascade *in vivo*. We reasoned that the afore-described metabolism-dependent antiviral mechanism of Propranolol may bypass cardiac side effects based on β-adrenergic receptor blockade and broaden the potential anti-coronavirus application of the widely-prescribed medicine. Interestingly, Propranolol is manufactured and prescribed as an enantiomeric mixture of two chiral forms, with the S-enantiomer exhibits ∼100 times more efficacy towards the β-adrenergic receptors than the R-enantiomer (*47, 48*). By contrast, both chiral forms displayed similar antiviral effects against βCoV in hepatocytes compared to the enantiomeric drug, as did the 50%-50% mixture of R- and S-forms (Fig. 5A, two enantiomers shown in S5A&B). Similar chirality-independent inhibitory effects on Alkyne-PC production of Propranolol enantiomers were also observed (Fig. S5C, quantified in S5D), further supporting a Lipin and metabolism dependent mechanism that could be safely explored to combat coronaviruses.

**Figure 5.**
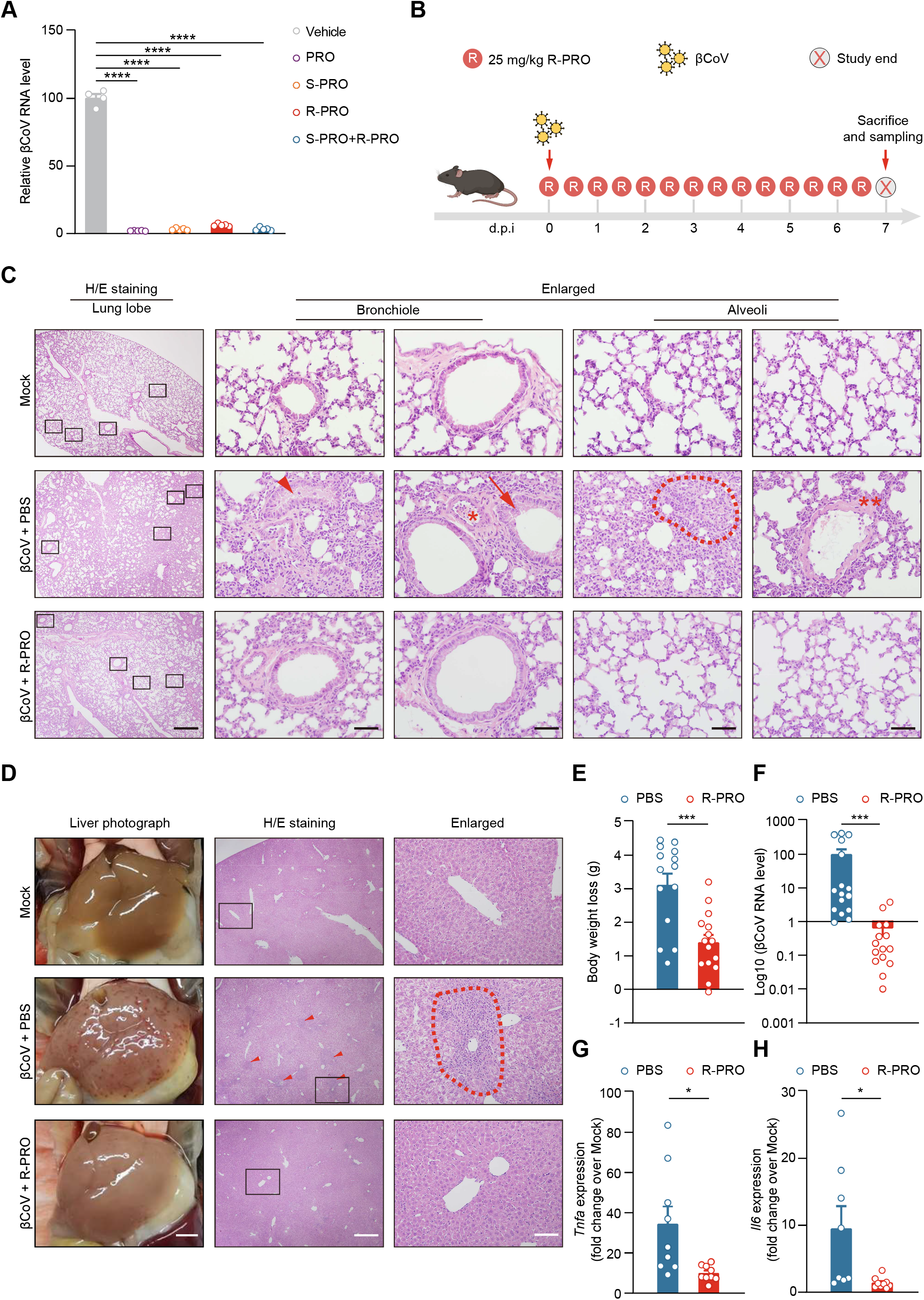
Propranolol exhibits broad protection against coronavirus pathology in mice. (A) Chirality-independent inhibition of coronavirus by Propranolol. Hepatocytes receiving 50 μM Propranolol or indicated R/S chirality are infected with βCoV, prior to RNA harvest for RT-qPCR. Representative results from 3 independent experiments are shown. Data are presented as mean ± SEM. ****, p < 0.0001. (B) Schematic depiction of Propranolol treatment in mice. βCoV-infected mice are treated with 25 mg/kg R-(+)-Propranolol or PBS every 12 hours for 7 days upon infection. (C) Propranolol treatment alleviates βCoV-induced pneumonia in mice revealed by H/E staining. βCoV infection in control mice causes lung bronchiole congestion (arrowhead), capillary hemorrhage (asterisk), bronchiole desquamation (arrow), and alveolar wall thickening with macrophages and lymphocytes infiltration (dash circle), hyaline membrane deposition (double asterisks), all of which are ameliorated by the Propranolol treatment. Representative data from 3 independent experiments are shown. Scale bars = 500 μm (left panels) or 50 μm (the rest panels). (D) Propranolol treatment alleviates βCoV-induced hepatitis in mice. βCoV infection in control mice caused wide spread liver damages, which are ameliorated by the Propranolol treatment. Red arrowhead and dash circles: inflammatory leukocytes surrounding necrotic foci. Representative data from 3 independent experiments are shown. Scale bars, 2.5 mm (left), 500 μm (middle), 100 μm (right). (E) Propranolol treatment decreases βCoV-induced body weight loss in mice. Body weight loss after 7 days post infection of untreated control (n = 14) or Propranolol-treated mice (n = 15) are shown. Representative data from 3 independent experiments are shown. Data are presented as mean ± SEM. ***, p < 0.001. (F) Propranolol treatment decreases βCoV viral loads in mice. βCoV N transcripts in the liver from untreated control (n = 10) or Propranolol-treated mice (n = 10) are determined by qPCR. Representative data from 3 independent experiments are shown. Data are presented as mean ± SEM. ***, p < 0.001. (G) Propranolol treatment decreases βCoV-induced inflammatory gene expression in mice. *Tnfa* transcripts in the liver from untreated control (n = 8) or Propranolol-treated mice (n = 8) are determined by qPCR. Representative data from 3 independent experiments are shown. Data are presented as mean ± SEM. *, p < 0.05. (H) *Il6* transcript levels are determined as in (G). Data are presented as mean ± SEM. *, p < 0.05.

The chiral-independent and broad-spectrum antiviral mechanism by Propranolol thus allowed us to employ the *in vivo* model with βCoV (MHV-A59) infection in mice (*8, 49, 50*), which represents an authentic form of coronavirus pathogenesis. This model also has the benefit of recapitulating the multiple organ pathologies seen with COVID-19 (*8, 49, 50*), and allows accurate delivery of substances/drugs via the intraperitoneal routes. Pilot experiments in a treatment regimen revealed that R-Propranolol displayed better efficacy against coronaviruses in mice compared to the S- or the enantiomeric form, likely reflecting cardiovascular effects of S-Propranolol (Fig. S6A&B). Therefore, we focused on the R-form in the treatment regimen, in which a dose of 25mg/kg R-Propranolol (drug dose/body weight) was given to the animal every 12 hours for 7 days as the animals were infected with coronaviruses (Fig. 5B). When converted with respect to drug metabolism rates based on body surface areas (BSA) in different species (*51*), the human equivalent dose (HED) of the above mice experiments is estimated to be ∼2mg/kg (drug dose/body weight), which can be tolerated in humans in general. Compared to non-infected mice, coronavirus infection caused acute tissue damages in the lung of saline-treated mice after 7 days as revealed by histology analysis (Fig. 5C, middle panels), including desquamation of the mucous layer of bronchioles and thickening of alveoli septa. The thickened alveoli walls were populated by increased number of type 2 pneumocytes and often infiltrated inflammatory cells, often with deposition of hyaline membranes that also indicated pulmonary injuries. Collectively, these pathologies generally resembled acute pneumonia and alveoli damages manifested in coronavirus diseases including those caused by COVID-19 (*8*). Propranolol largely protected the animals from the viral-induced pneumonia (Fig. 5C, lower panels). Moreover, murine coronavirus also caused drastic liver damages in PBS-treated mice (Fig. 5D, middle panels), as evidenced by gross morphology or histological analysis. Propranolol treatment also alleviated the hepatitis caused coronavirus infection in mice (Fig. 5D, lower panels). As a result, Propranolol treatment reduced weight loss caused by coronavirus infection (Fig. 5E).

Consistent with the pathological analysis, viral loads were decreased by ∼99.7% in the liver of treated mice compared to untreated ones (Fig. 5F). The induction of inflammatory factors such as TNFα or IL6 was also blunted (Fig. 5G&H). These data support the therapeutic potential of R-Propranolol in countering diseases caused by coronavirus infection that is independent of its original use as beta-blockers (*48, 52*).

### Moderate dose of Propranolol as anti-coronaviral prophylactics

Although several antivirals have been approved to treat coronavirus diseases such as COVID-19, there is no approved drug for prevention to date, likely due to higher safety bars in prophylactic medicines. Propranolol has been widely prescribed, readily supplied and generally well tolerated (*39, 53*). Moreover, multiple new versions of beta-blockers have been developed and are extensively used for cardiovascular complications, yet apparently without considering Lipin inhibition. This discrepancy led us to perform a functional survey on different beta-blockers for their “moonlight” anti-coronavirus function mostly likely via targeting Lipin1. Interestingly, the different beta-blockers showed a wide range of anti-coronavirus effects in cells, with Propranolol, Nebivolol, and Penbutolol being most effective (Fig. 6A). The antiviral efficacy of these beta-blockers appeared to depend on their lipophilic nature (Fig. 6B), correlating with the lipogenic role of Lipin as well as the molecular docking between Propranolol and Lipin1 (Fig. 6C).

**Figure 6.**
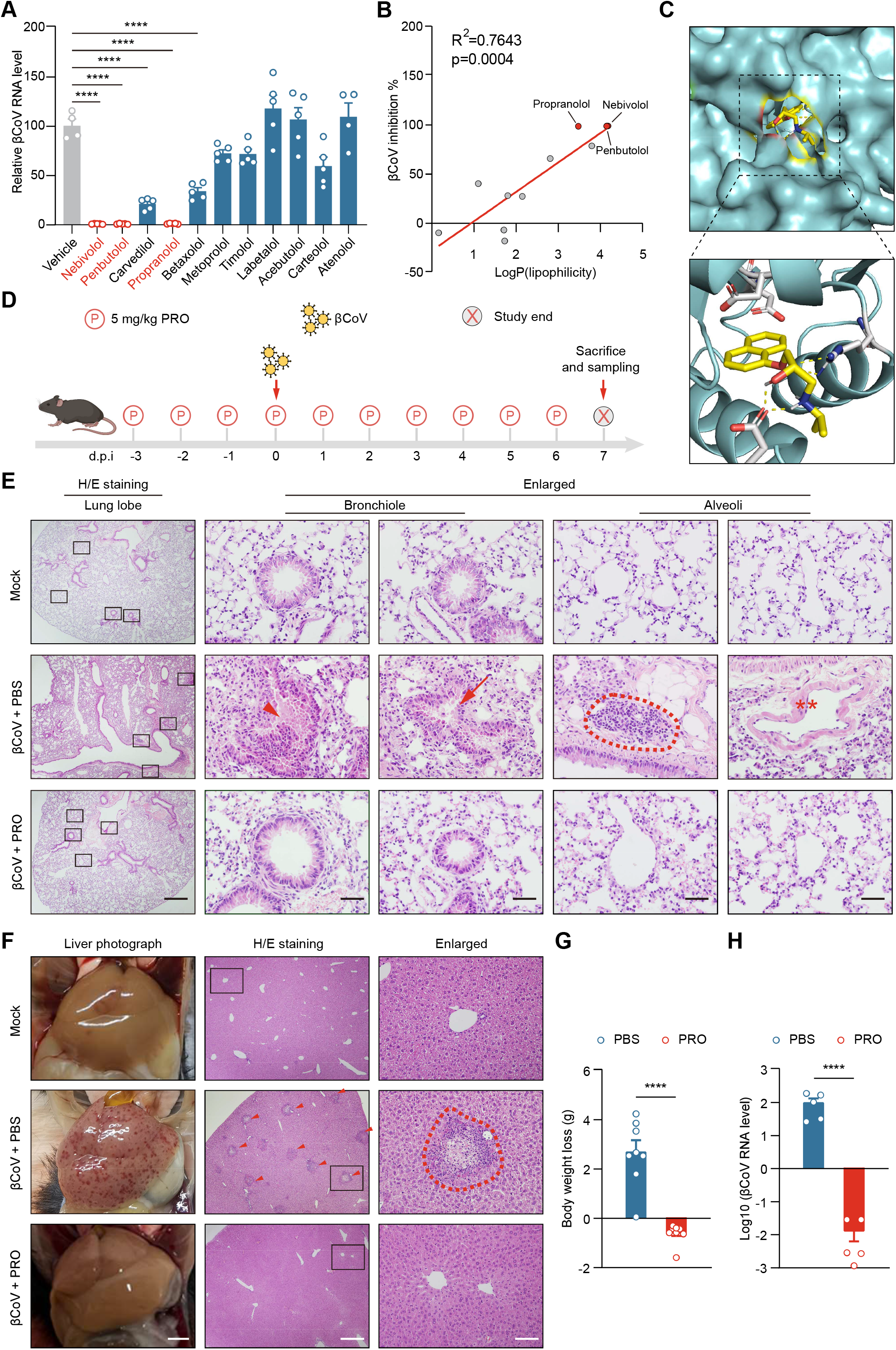
Prophylactic application of Propranolol against coronavirus in mice. (A) Different beta-blockers show a wide range of anti-coronavirus effects. Primary hepatocytes pretreated with 50 μM beta-blockers for 12 h were infected with βCoV for 8 h, prior to harvested for RT-qPCR. Data are presented as mean ± SEM. ****, p < 0.0001. (B) The correlation analysis of beta-blockers βCoV inhibition ability and their lipophilicity. (C) The molecular docking between Propranolol and the catalytic domain of human Lipin1. (D) Schematic depiction of the prophylactic application of Propranolol in mice. βCoV-infected mice are treated with 5 mg/kg Propranolol or PBS every day, starting 3 days before infection. Mice are sacrificed after 7 days post infection. (E) Propranolol prevents βCoV-induced pneumonia in mice revealed by H/E staining. βCoV infection in control mice causes lung hemorrhage and capillary congestion (arrow head), bronchiole desquamation (arrow), alveolar wall thickening with macrophages and lymphocytes infiltration (dash circle) and hyaline membrane deposition (double asterisks), all of which are ameliorated by the Propranolol treatment. Representative data from 3 independent experiments are shown. Scale bars = 500 μm (left panels) or 50 μm (the rest panels). (F) Propranolol prevents βCoV-induced hepatitis in mice. βCoV infection in control mice caused wide spread liver damages, which are ameliorated by the Propranolol treatment. Red arrowhead and dash circles: inflammatory leukocytes surrounding necrotic foci. Representative data from 3 independent experiments are shown. Scale bars, 2.5 mm (left), 500 μm (middle), 100 μm (right). (G) Propranolol prevents βCoV-induced body weight loss in mice. Body weight loss after 7 days post infection of untreated control (n = 8) or Propranolol-treated mice (n = 8) are shown. Representative data from 3 independent experiments are shown. Data are presented as mean ± SEM. ****, p < 0.0001. (H) Propranolol diminishes βCoV viral loads in mice. βCoV N transcripts in the liver from untreated control (n = 5) or Propranolol-treated mice (n = 5) are determined by qPCR. Representative data from 3 independent experiments are shown. Data are presented as mean ± SEM. ****, p < 0.0001.

The wide prescription of Propranolol and related medicines intrigued us to examine preventative effects of Propranolol against coronavirus infection, particularly since its host-based antiviral mechanism may enable a broad-spectrum potential for countering the constantly-mutating coronavirus. To this end, a 3-day, moderate dose (5mg/kg/day) pre-treatment of Propranolol was combined with daily treatment for 7 days after coronavirus infection (Fig. 6D). This dose is within the range of prescription to adults for cardiovascular complications, as well as of that could be prescribed to infants for treating hemangioma (*43, 44*).

Of note, the preemptive strategy provided significant protection from pathologies in the lung caused by coronavirus infection (Fig. 6E, middle vs lower panels), largely eliminating the otherwise severe tissue damages. Similar protection on the liver was also observed (Fig. 6F). Accordingly, mice pretreated with a moderate dose of Propranolol displayed little if any weight loss compared to βCoV-infected controls (Fig. 6G), indicating an overall protection at the systemic level. On the molecular level, the drug drastically reduced viral load by over 4 orders compared to βCoV-infected animals without treatment (Fig. 6H). Moreover, IB on viral proteins or qPCR of viral genes showed that the preventative measure reduced viral loads close to the background levels in non-infected mice (Fig. S7A). Similar prevention of inflammatory gene induction such as *Tnfa* or *Il6* was also observed (Fig. S7B&C). Together with the mechanistic studies, the data support the *in vivo* efficacy of the repurposed drug in preventing coronavirus infection, providing a metabolism-oriented strategy for developing host-based medications as safe and broad-spectrum antivirals to combat the current and future coronavirus diseases.

## Discussion

The high mutation rates, combined with high infectivity and stability, of zoonotic coronaviruses pose great challenges for developing effective antivirals (*3*). Besides inhibiting viral factors, targeting the host pathways required for the viral life cycle may therefore provide a parallel strategy for antivirals, with the potential of being broad-spectrum (*54*). Moreover, mutation rates for host genes are thought to be much lower than viral genes (*6, 7*), thereby slowing down the emergence of drug resistance. In this regard, our strategy of identifying the druggable VSE factors and the subsequent metabolic targeting may achieve both antiviral efficacy and host safety, therefore deserving further attention. Such factors may be identified through mining cellular networks of known host factors, or dissecting viral-specific events in the host. The replication organelle, a unique yet essential host structure induced by the coronavirus (*6, 11*), may represent an effective and plausible target to counter the pathogen. One may even speculate that limiting the membranous RO formation by Propranolol, as tested in the current study, could disrupt the spatial assembly of viral replication complexes and increase the accessibility of viral-based drugs for targeting these functional machineries. In principle, this combination strategy could alleviate the shortage of medicine, reduce the cost of therapy, and likely decrease the risk of the emergence of drug resistance.

The continuing unraveling of coronaviruses life cycle has in turn advanced our understanding of the biology in host cells (*6, 7*). As shown in the fundamental process referred as membragenesis, the master regulator SREBP1 concertedly turns on both the production of phospholipids (via metabolic enzymes such as Lipin) and the assembly of bilayer (via the TMEM41B scramblase). Coronaviruses therefore may particularly rely on the membragenic mechanism to boost the construction of membranous replication compartments and likely the subsequent assembly of virion envelopes (*11, 14*), a mechanism shared by other RNA virus (*11, 13, 55*). Moreover, given the universal demand of increased membrane supply with bilayer equilibration in numerous processes such as cell growth, division, differentiation, metabolism, etc., however, the SREBP1-TMEM41B regulatory circuit may emerge as a key “rheostatic” mechanism that deserves further investigation and warrants relevance to both cells and viruses. Consistent with the notion, the cytosolic enzyme Lipin represents a central and highly regulated node in glycerophospholipid production, and has been implicated in metabolism, cancer, as well as tissue homeostasis (*36, 56, 57*). Our finding that Propranolol could be repurposed as a potential broad-spectrum anti-coronavirus agent, though serendipitous, is indeed driven by the understanding of host lipid metabolism (*37*) and particularly SREBP1-TMEM41B in membragenesis. One may foresee that additional specific inhibitors could be developed to limit the metabolic fluxes of phospholipids to target this pivotal host pathway.

Although our current study is limited to cellular and murine models of coronavirus pathogenesis, the work could be of broad clinical relevance, inasmuch that Propranolol has been once wildly prescribed with extensive sets of pharmacology and toxicology data in humans (*39, 53*). Propranolol is generally safe, orally active, and available in generic forms. While the drug is commonly prescribed as a 50-50% enantiomeric mixture with the S-form being more relevant to cardiovascular complications (*47, 48*), we found that both chiral forms effectively inhibit coronaviruses and R-propranolol may offer better therapeutic outcomes *in vivo*. Hence, different combination of the two chiral forms could allow additional leverage of Propranolol’s beneficial effects on cardiovascular diseases (*52*) that often exacerbate COVID-19, thereby further reducing mortalities and helping recovery from the disease. These additional indications, and most importantly the anti-coronaviral effects by Propranolol in patients in the treatment or particularly the prophylactic regimens, await support by real world data and/or demonstration by clinical trials.

## Acknowledgements

The authors thank Drs. R. Schekman (UC Berkely), J. Liu (Mayo Clinic), and D. Ginsburg (Michigan) for helpful discussions and critical reading of the manuscript. In addition, the authors thank Dr B. Song (Wuhan University) for providing *SCAP*^*flox/flox*^ mice and Dr N. Tang (National Institute of Biological Sciences, Beijing) for A549 cell line.

## Author Contributions

Conceptualization: X.W.-C.;

Methodology: Z.Q.;

Investigation: H.F., L.L., Y.W., K.C., P. L., X.X., Y.Y., F.Z., L.W., Y.Z., B.X., D.H.,Z.Q. and X.W.-C.;

Writing: X.W.-C.;

Supervision: X.W.-C. and Z.Q.

## Data and materials availability

Further information and requests for reagents and resource will be fulfilled by the lead contact, Xiao-Wei Chen.

## Supplementary Materials

Materials and Methods

Reference (1)

Movies S1 to S2

**Supplementary Figure 1.**
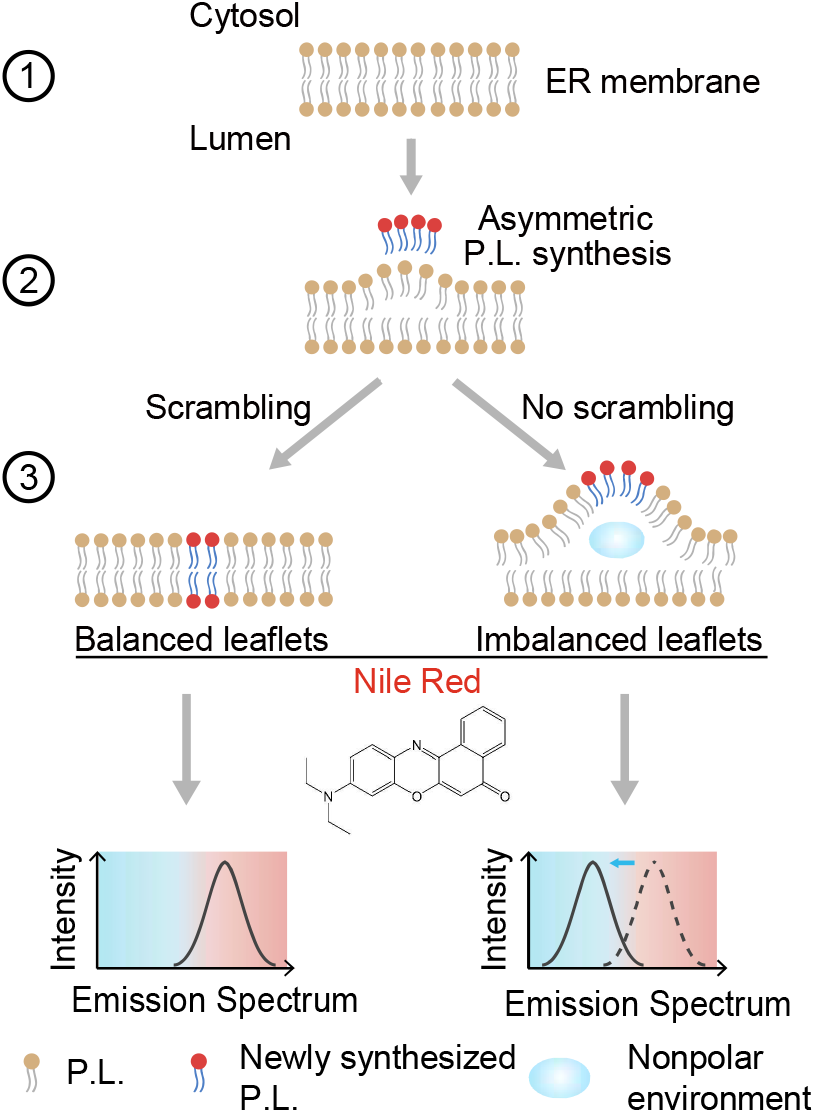
Ratiometric imaging of membrane environment with the solvatochromic dye Nile Red. Schematic diagram of imbalanced phospholipid addition to the ER bilayer during membrane biogenesis. : existing ER bilayer. : asymmetric phospholipid addition at the cytosolic leaflet. : lipid scrambling to produce balanced bilayer (left), and imbalanced or damaged bilayer caused by deficient lipid scrambling (right). Lower: The solvatochromic lipid dye Nile Red detects the lipid environment alterations upon scrambling defect and exhibits changes in the emission spectrum.

**Supplementary Figure 2.**
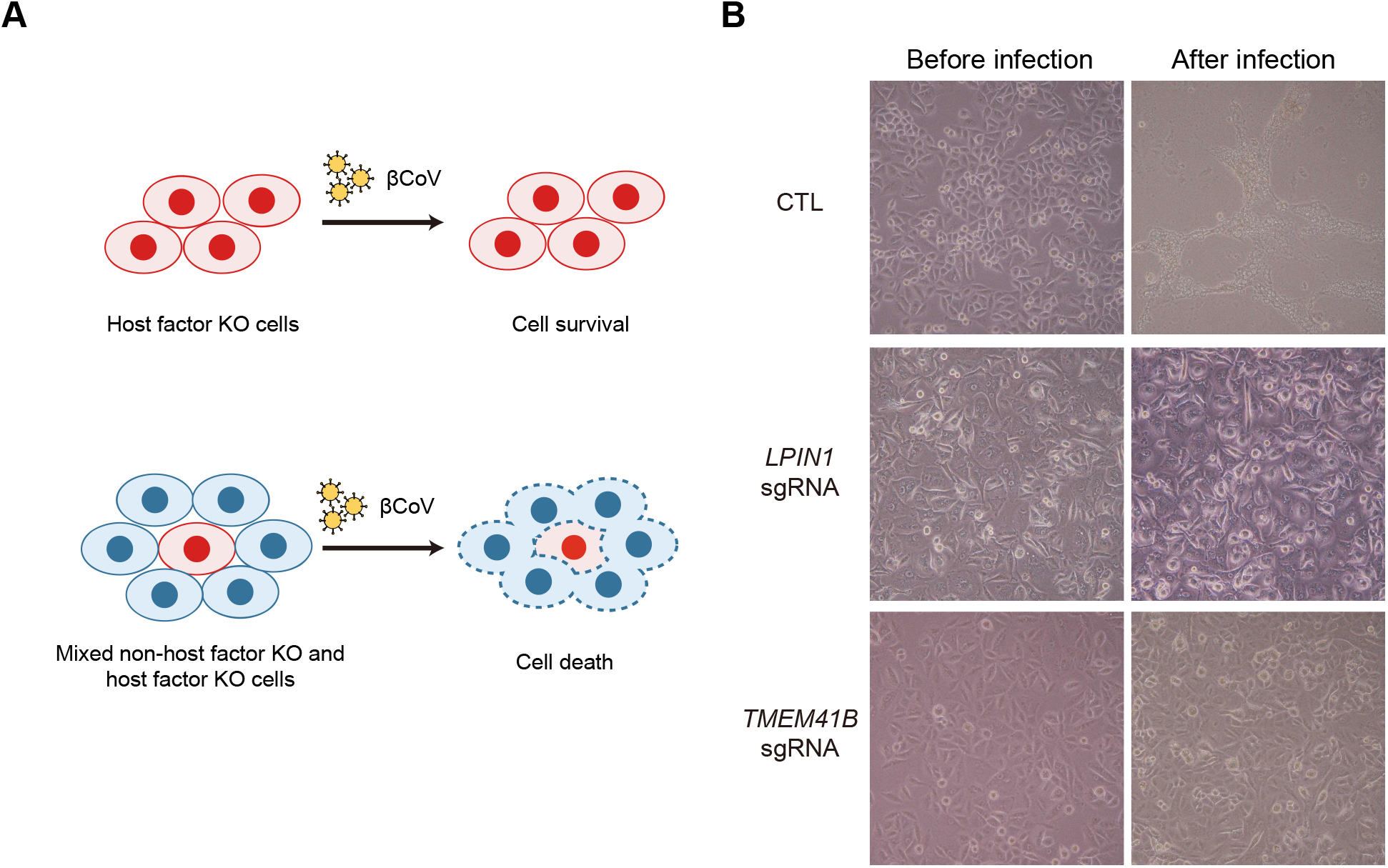
*LPIN1* ablation inhibits βCoV-induced syncytia formation and the virtual screen results. (A) Scheme of the arrayed CRISPR screen which identifies host factors that may be missed in the pooled CRISPR screen. (B) Microscopy images of wild type, LPIN1 or TMEM41B knockout A549mCEACAM1 cells before or after βCoV infection for 48 hours.

**Supplementary Figure 3.**
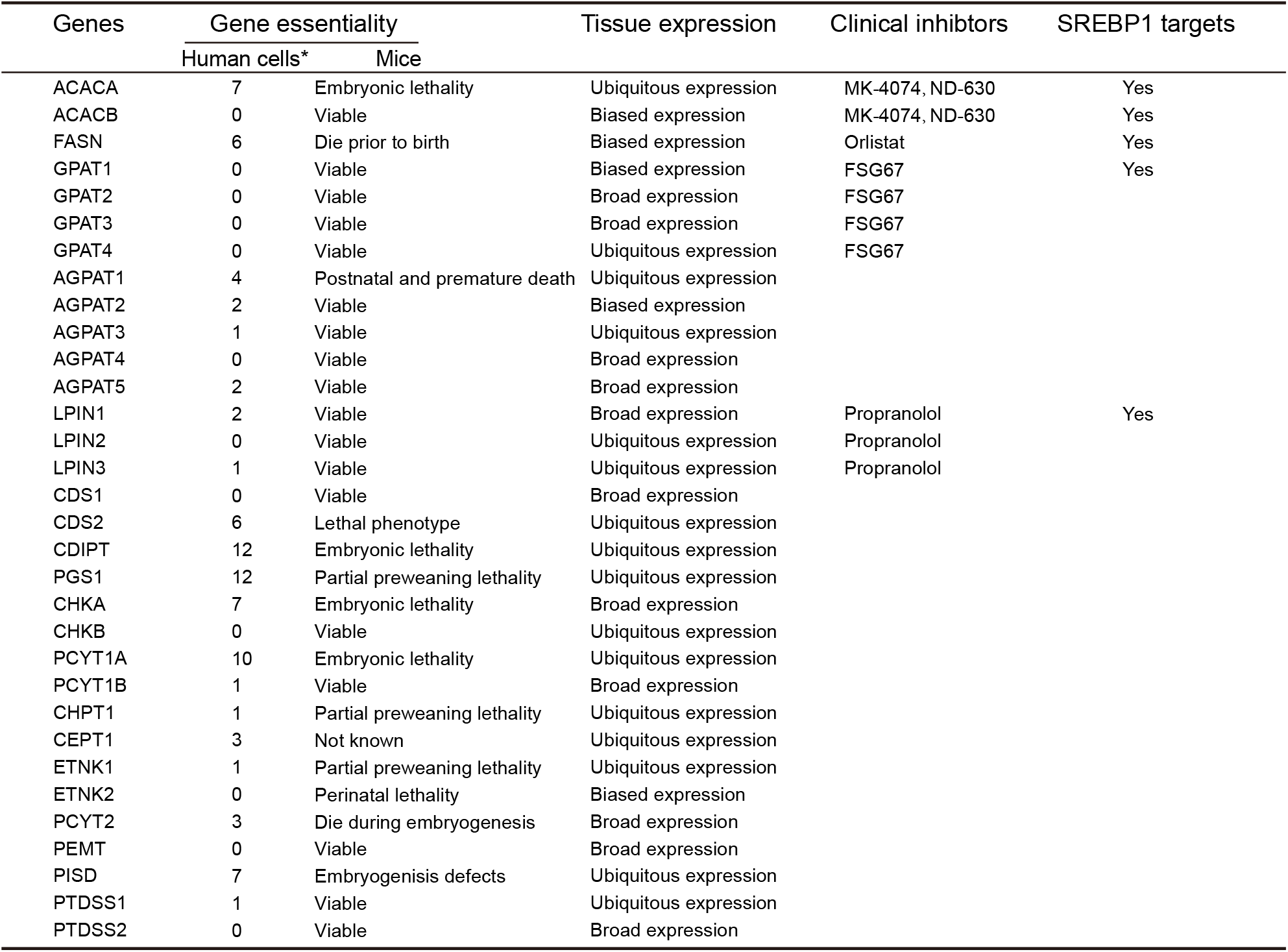
Virtual analysis of genes related to phospholipid biosynthetic pathway. *The numbers refer to how many studies reported the gene essentiality in human cells from the database (http://origin.tubic.org/deg), three as the threshold.

**Supplementary Figure 4.**
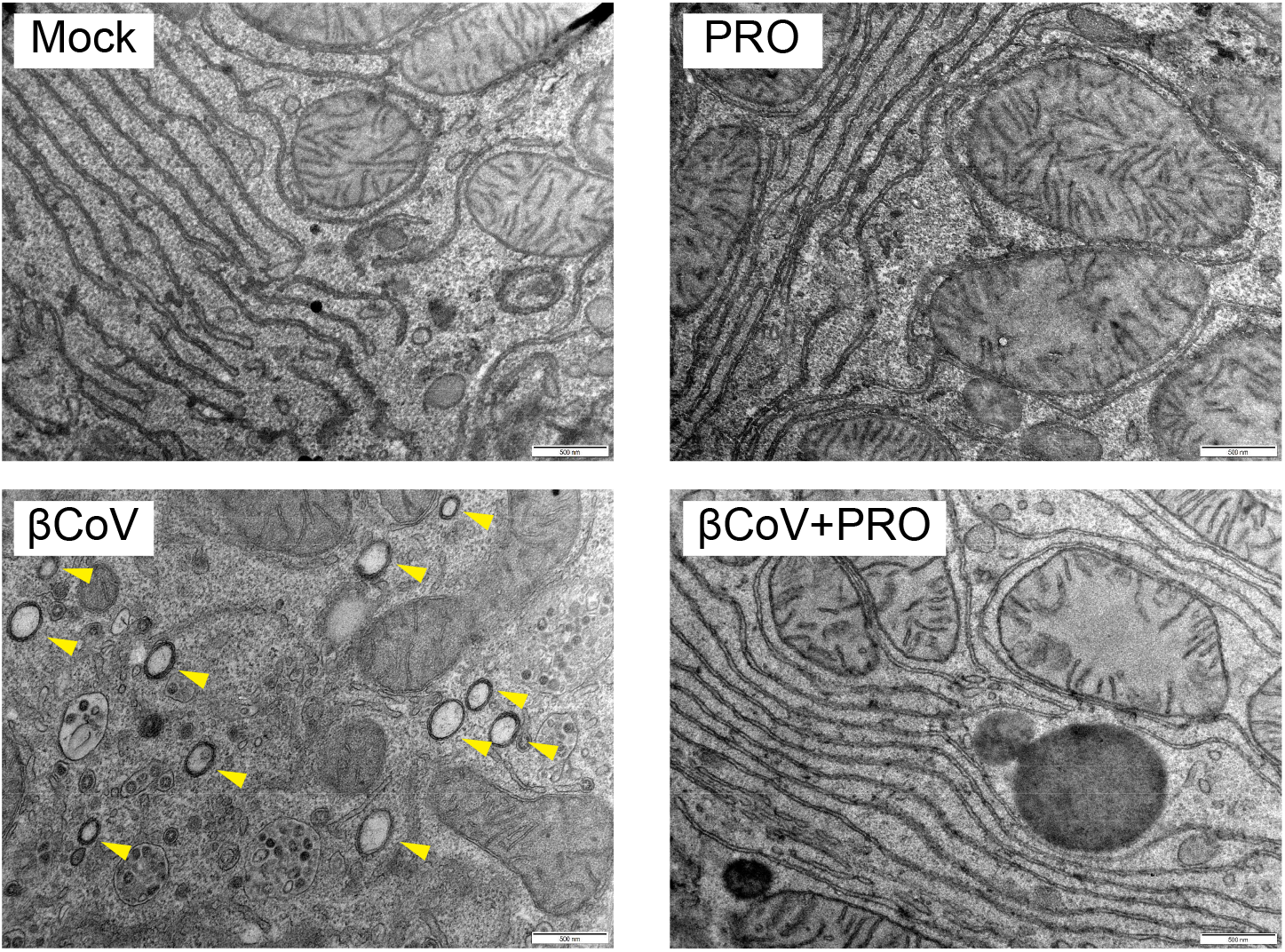
βCoV induced replication organelles eliminated by Propranolol in hepatocytes. Representative TEM images of the replication organelles (ROs) induced by βCoV and their elimination by Propranolol. Primary hepatocyte samples were fixed and subjected to ultra-thin section, followed by TEM analysis. Yellow arrowheads indicate the double membrane vesicles (DMVs). Scale bars=500 nm.

**Supplementary Figure 5.**
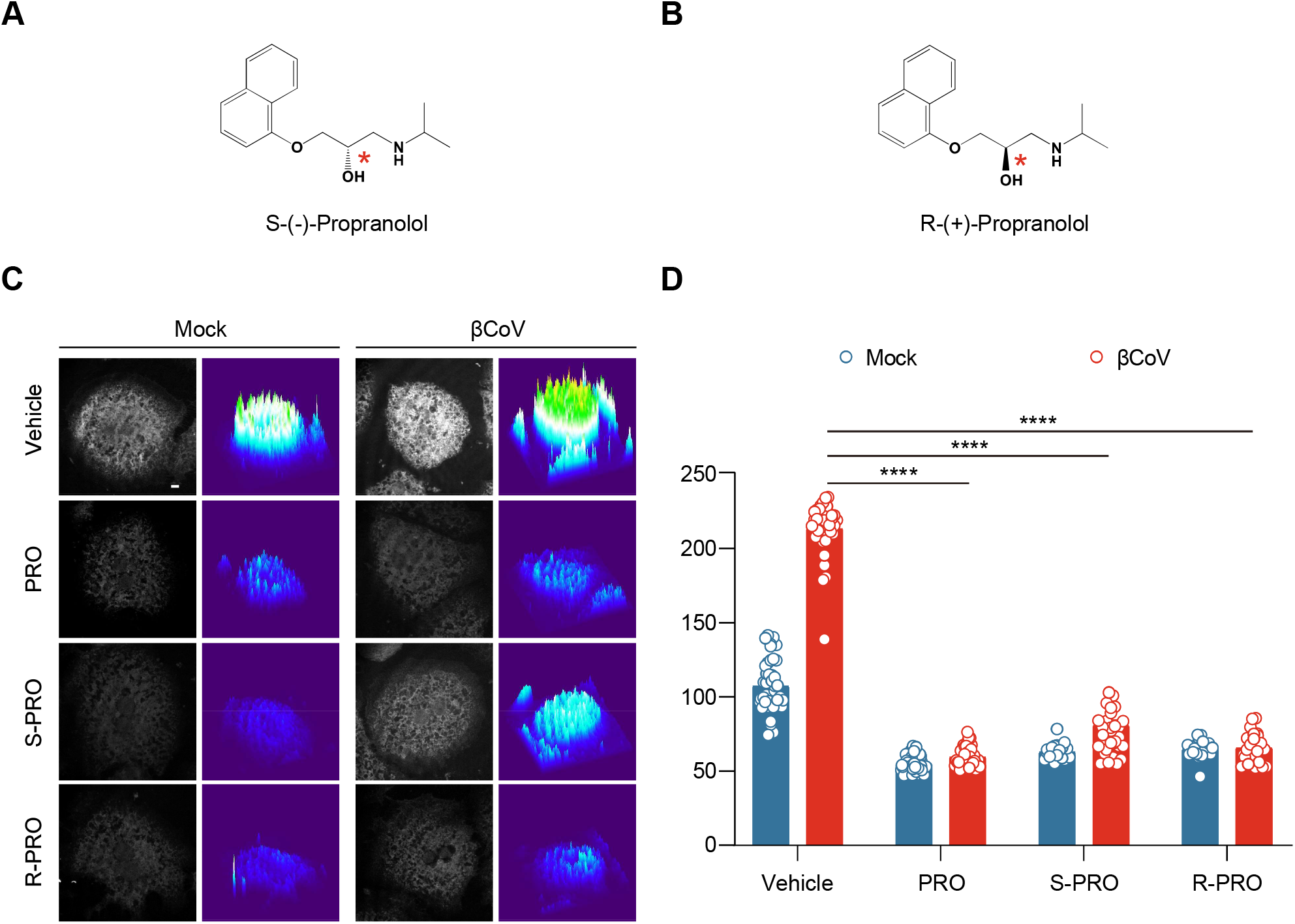
S(-)-Propranolol and R-(+)-Propranolol exhibit similar βCoV induced phospholipid synthesis lowering effect. (A) Chemical structure of the S-(-)-Propranolol. Dashed line indicates that the bond is extending behind the plane of the drawing. The chiral center is highlighted by the red asterisk. (B) Chemical structure of the R-(+)-Propranolol. Bold-wedged line indicates that the bond is protruding out from the plane of the drawing surface. The chiral center is highlighted by the red asterisk. (C) S-(-)-Propranolol and R-(+)-Propranolol decreases alkyne-PC production induced by βCoV infection. Primary hepatocytes receiving vehicle or 50 μM S-(-)-Propranolol or R-(+)-Propranolol are uninfected or infected with βCoV. Cells are labeled by alkyne-choline prior to click conjugation and confocal microscopy. Representative results from 3 independent experiments are shown. Left: confocal images. Right: surface plots. Scale bar=5 μm. (D) Quantification of alkyne-PC signals from (C). Data are presented as mean ± SEM. ****, p < 0.0001.

**Supplementary Figure 6.**
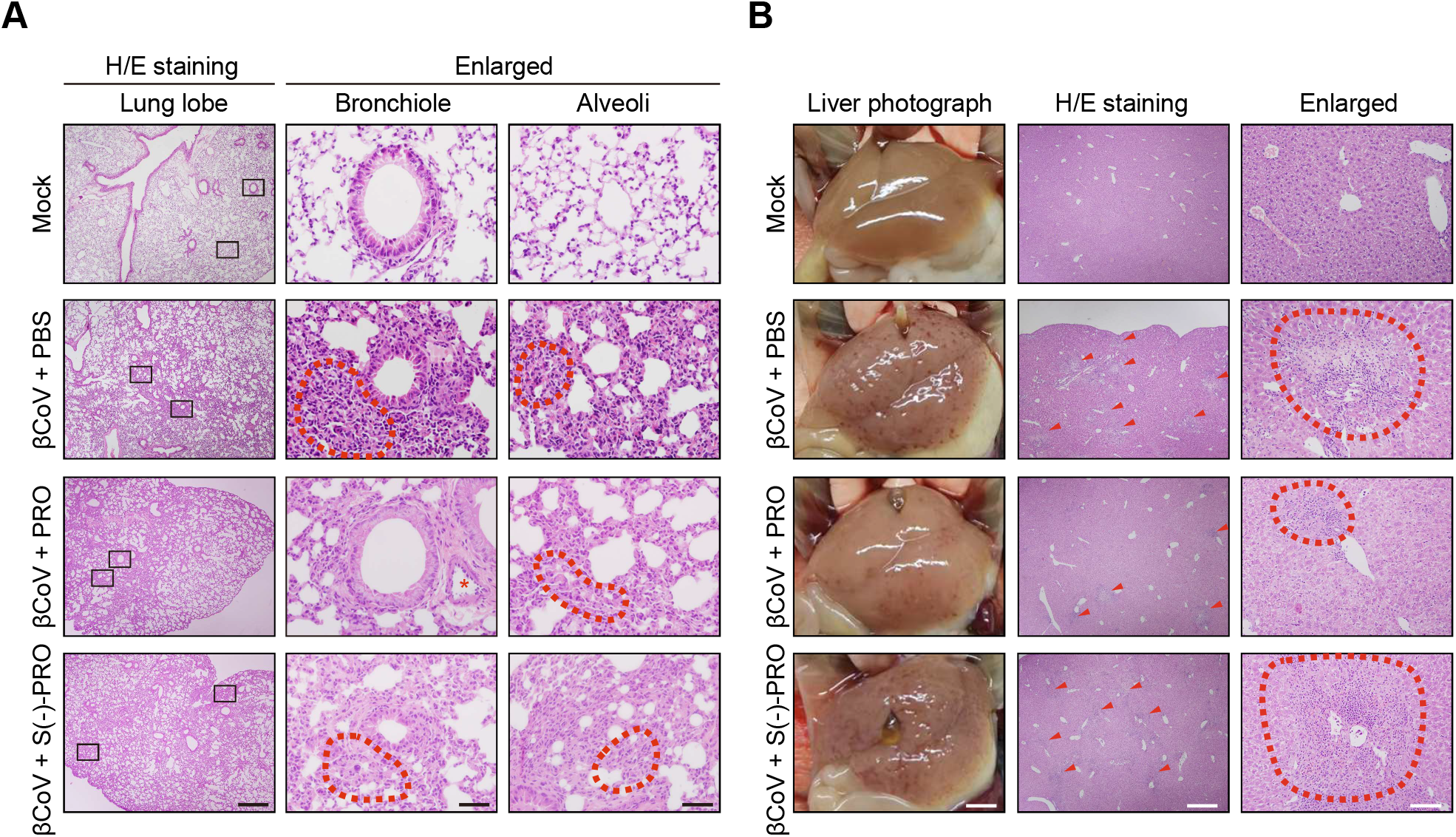
Pilot therapeutic experiments with enantiomeric and S-(-)-Propranolol in mice. (A) S-(-)-Propranolol exhibits little protection from βCoV-induced pneumonia in mice revealed by H/E staining as in Figure 4. The βCoV infected mice exhibits alveolar wall thickening with macrophage and leukocyte infiltration (dash circle) which are ameliorated by the treatment with the enantiomeric Propranolol, while little alleviation is observed in the lung of the S-(-)-Propranolol treated mice. Black rectangles indicate the enlarged viewed areas. Scale bars=500 μm (left panels) or 50 μm (the rest panels). (B) S-(-)-Propranolol exhibits little protection βCoV-induced hepatitis in mice. βCoV infection in un-treated mice induced widely distributed liver damages which are ameliorated by the treatment with the enantiomeric Propranolol, whereas little amelioration shown in the liver from the S-(-)-Propranolol treated mice. Black rectangles indicate the enlarged viewed areas. The leukocytes and macrophages surrounding necrotic foci are highlighted by the red arrowhead and dash circles. Scale bars, 2.5 mm (left), 500 μm (middle), 100 μm (right).

**Supplementary Figure 7.**
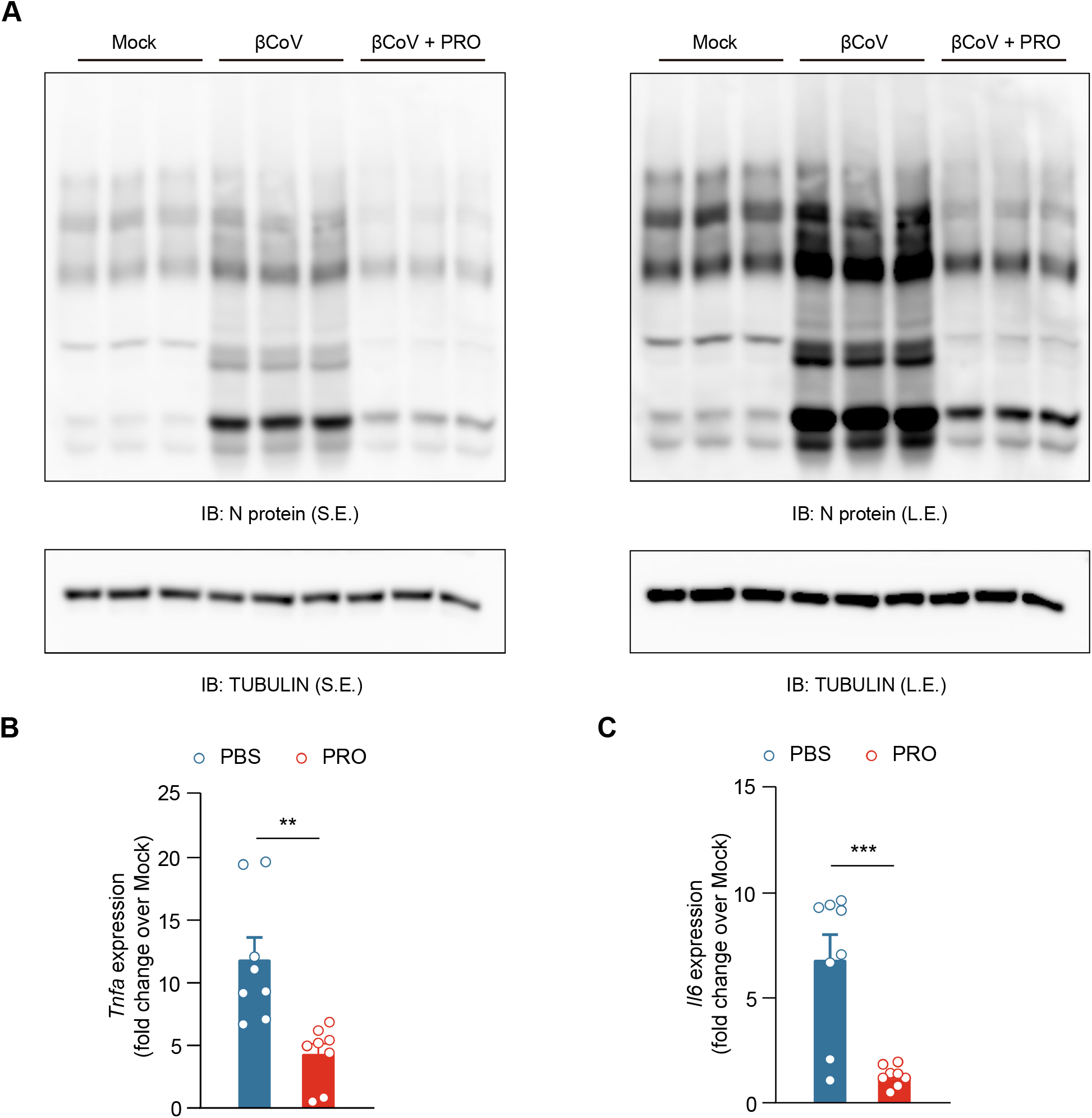
Propranolol’s preventative measure reduces βCoV infection induced viral protein expression and inflammatory gene induction. (A) βCoV infection induced viral protein expression is diminished by the Prorpanolol prophylactic application revealed by IB. Liver lysates of uninfected (Mock), βCoV infected (βCoV) and Propranolol treated βCoV infected mice (βCoV+PRO) as in Figure 6 are used for IB with an anti-N protein antibody. TUBULIN is used as an internal control. Left: S.E., shorter exposure. Right: L.E., longer exposure. (B) Propranolol prevents βCoV-induced inflammatory gene expression in mice. Tnfa transcripts in the liver from untreated control (n = 8) or Propranolol-treated mice (n = 8) are determined by qPCR. Representative data from 3 independent experiments are shown. Data are presented as mean ± SEM. **, p < 0.01. (C) Il6 transcript levels are determined as in (B). Data are presented as mean ± SEM. ***, p < 0.001.

