## Supplementary material for "A Host-Harbored Metabolic Susceptibility of Coronavirus Enables Broad-Spectrum Targeting": Materials and methods

**Supplementary materials**

**Materials and Methods**

**Mouse models**

Animal housing and all the experimental procedures are authorized by the Institutional Animal Care and Use Committees of Peking University. All mice used for experiments are on C57BL/6 background. Un-infected animals are housed in the specific pathogen free (SPF) animal facility of the Peking University and the facility was certified by Association for Assessment and Accreditation of Laboratory Animal Care (AAALAC). Experiments with beta-coronavirus (Murine hepatitis virus-A59) infected mice are carried out in a P2 laboratory. All animals are housed with a 12 h light/dark cycle (lights on: 06:00-18:00). Distilled water and mouse chow diet are provided *ad libitum* unless otherwise noted. *SCAP^flox/flox^* mice are kindly provided by B. Song (Wuhan University). LoxP-Cre mediated spCas9 knockin (KI) mice are purchased from Jackson Lab (JAX: 026556, RRID:IMSR_JAX:026556) to generate hepatic *Lpin1* knockout and *Tmem41b* knockout as previously described (*1*). Wild type C57BL/6 mice are purchased from Charles River. All mice except those from Charles River are generated by in-housing mating. Mice from Charles River are imported into the facility prior to experiments.

**Adeno-Associated Virus (AAV) plasmid construction, production and delivery**

AAV plasmids are constructed by homologous recombination. pX602-TBG-Cre-sgRNA vector was generated from the pX602 (Addgene, 61593) vector. In brief, Cre recombinase is recombined into the AgeI and EcoRI sites, and the U6-target-scaffold fragment is recombined into the NotI and KpnI sites. The target sequence of sgRNA is designed by the Benchling platform and listed in supplementary table 2. Expression vector AAV-TBG-cDNA is generated from AAV-TBG-GFP (Addgene, 105535). cDNA is cloned from a cDNA library and then recombined into the NotI and BamHI sites. All clones are verified by sanger sequencing.

AAV is packaged by HEK293T cells and purified by density gradient centrifugation as previously described. In brief, HEK293T cells are transfected with AAV shuttle plasmids, Rep/Cap (2/8) plasmids, and helper plasmids by PEI. After 48 h post-transfection, cells are harvested for AAV purification using Opti-prep gradient. Purified AAV is delivered into mice through tail vein injection at 4E11 virus titer per mouse for CRISPR-mediated knockout in mice liver.

**Cell culture**

All cells are maintained in Dulbecco modified Eagle medium (DMEM) supplemented with 10% FBS and 1% P/S at 37°C incubator with 5% CO_2_. HEK293T (ATCC, CRL-3216, RRID:CVCL_0063) and Huh7 (JCRB Cell Ban, JCRB0403, RRID:CVCL_0336) cells are obtained from ATCC and JCRB, respectively.

Mouse primary hepatocytes are isolated from liver of male mice with indicated genotypes. *Lpin1* knockout and *SCAP^fl/fl^* mice primary hepatocytes are isolated from 10-week-old male mice after 4 weeks upon AAV mediated KO. *Tmem41b* KO mice primary hepatocytes were isolated from 10-weeek-old mice after 2 weeks upon AAV mediated KO. Mice are anesthetized by 1% Pelltobarbitalum Natricum (AMRESCO, USA) prior to dissection and perfusion first with 50 mL Krebs Ringer with Glucose (KRG) buffer (37℃) via the vena cava to remove the residual blood, followed by 30 mL type IV collagenase (C5138, sigma) in KRG buffer for cell dissociation. Thereafter, the liver is isolated from mice immediately, cut into pieces and filtered through a 70 μm cell strainer (352350, Falcon) to remove tissue debris. The digested livers are washed for 3 times with cold DMEM followed by pelleting at 50 g for 4 min. The isolated primary hepatocytes are maintained in DMEM medium with 10% FBS and 1% penicillin and streptomycin. Isolated primary hepatocytes are infected with MOI = 0.5 beta-coronavirus (MHV-A59) 12 h post isolation. The virus containing DMEM medium is replaced with DMEM medium with 10% FBS and 1% P/S after 1 h.

**MHV production and viral infection**

Beta-coronavirus (βCoV) mouse hepatitis virus A59 strain (MHV-A59) is provided by K. Holmes (University of Colorado). The virus stock is propagated in 17Cl-1 cells followed by 3 cycles of freeze and thaw. After a spinning down to clear cell debris, the supernatant is used as a working stock. The titer of the virus stock is determined by a plaque assay in 17Cl-1 cells. Conditions for βCoV infection in cells and mice are specified in later sections of experimental procedures.

**Arrayed CRISPR-Cas9 Screen**

A549 cells are generously provided by N. Tang (National Institute of Biological Sciences, Beijing). Cas9 and mCEACAM1 stably expressed A549 cells utilized for the CRISPR-Cas9 screen are constructed by lentivirus transduction. The arrayed sub-library of phospholipid metabolism gene sgRNA is acquired from Sanger Arrayed Whole Genome Lentiviral CRISPR Library Human, Glycerol Format (HSANGERG, Sigma-Aldrich). HEK293T cells are seeded at 60% confluence in a six well plate the day before transfection. Each well is transfected with 2 μg sgRNA plasmid and lentiviral package plasmids mixed in PEI contained culture medium. The sgRNA packaged lentivirus is harvested at 48 hours after transfection. The A549-Cas9-mCEACAM1 cells at 60% confluence are infected with sgRNA packaged lentivirus for 48 hours prior to puromycin selection for 4 days. Then A549 cells are infected with βCoV at MOI = 0.001. After 48 hours infection, the images of cell fusion and death are acquired by a light microscopy.

**Luciferase reporter assay**

For the human active SREBP1 binds to the SRE in TMEM41B promoter experiment. HEK293A cells cultured in 6-well plates are transfected with luciferase reporter together with Renilla, and active SREBP1 or GFP using Lipofectamine 2000 (Invitrogen). For SARS-CoV-2 replicon experiment, HEK293T cells cultured in 6-well plates are pretreated with Propranolol for 6 h prior to transfection, and transfected with a SARS-CoV-2 replicon plasmid using Lipofectamine 2000 (Invitrogen) according to the manufacturer’s protocol. Cells are collected at 36 h post transfection, by rinsing with cold PBS once after medium removal, and lysed with 1 x passive lysis buffer (Promega, E1910) for 15 min. Luciferase activity in cell lysates is measured by the luciferase reporter assay system (Perkin Elmer, United Kingdom) according to the manufacturer’s instructions.

**Immunoblot (IB)**

Whole-cell lysate (WCL) is extracted using cell lysis buffer (150 mM NaCl, 50 mM Tris pH 7.5, 1 mM EDTA, 1% SDS, 10% glycerol, and protease inhibitor). WCL is centrifuged at 10,000 rpm for 10 min at 4℃, and the supernatants are collected and mixed with sample buffer prior to by 3-15% Tris-acetate SDS-PAGE. The proteins are transferred to the nitrocellulose filter membrane and incubated with specific primary antibodies and HRP-conjugated secondary antibodies before visualization by ECL.

**Immunofluorescence microscopy**

Cultured cells or primary hepatocytes plated on glass coverslips are washed with PBS twice before methanol fixation. Then the cells are rehydrated in PBS and blocked by PBS solution containing 2% goat serum for 20 min. After incubating with primary antibodies overnight at 4℃, the cells are washed with PBS three times and then incubated with the second antibodies for 2 h before imaging. Immunostaining images are collected in a double-blind manner using Zeiss spinning disk microscopy and quantified using ImageJ (Fiji).

**Ratiometric imaging with the solvatochromic Nile Red dye**

For the living cell imaging, the cells are seeded onto a chambered coverslip at a density of 5 × 10^4^ cells/well 24 h before infection. The cells are infected by AAV for 24 h or by MHV-A59 for 16 h before imaging. For labelling, 1 µg/mL Nile Red (N1142, Invitrogen, USA) is added into the culture medium 10 min before imaging. Ratiometric imaging is performed using a Live SR CSU W1 microscope, equipped with 100X oil (NA = 1.4) objective, Live SR Structured Illumination Super Resolution module. The excitation light is provided by a 488 nm laser, while the emission fluorescence is collected at two spectral ranges: 580-653 nm (I580-653) and 665-705 nm (I665−705) in sequential mode by rapid switching to minimize drift. The E.I. (Emission Index) is calculated by dividing the I580−653 channel image by the I665-705 channel image. MATLAB (MathWorks, Natick, MA) is used to calculate the two-dimensional (2D) E.I. map.

**Click labeling with alkyne-choline**

Primary hepatocytes are seeded on glass coverslips in 6-well plates at about 50% confluence. Primary hepatocytes are pretreated with 50 μM propranolol for 12 h, and then labelled with 100 μM alkyne-choline for another 4 h. For infection experiments, the cells are infected with MHV-A59 for 8 h before labeling. The cells are washed by warm PBS (37℃) twice, fixed by 4% (w/v) formaldehyde for 15 min, rehydrated in PBS for 15 min, and finally permeabilized by 25 μg/mL digitonin for 10 min. The cells on coverslips are incubated with click reaction solution (10 μM 5-TAMRA azide, 50 μM BTTAA-CuSO4 complex (BTTAA/CuSO4 6:1, mol/mol) and 2.5 mM sodium ascorbate) for 1 h at RT. After click-reaction, cells are mounted and subjected to confocal microscopy in a double blinder manner and the signals are quantified using ImageJ (Fiji).

**Liposomes preparation and delivery**

Phospholipids are purchased from Avanti Polar Lipids and dissolved in chloroform. ER-targeting liposomes are generated as described (*2*) and are composed of DOPC/DOPE/DOPS at a molar ratio of 2: 2: 1. For liposome rescue experiments, primary hepatocytes are infected with MHV-A59 for 2 h, and then the culture medium is replaced by liposomes containing medium with a final phospholipid concentration of 60 μM. Cells are harvested for RNA extraction after 6-h incubation.

**Electron microscopy and FIB-SEM**

Primary hepatocytes cultured on polyvinylchloride (PVC) membrane are washed with PBS twice before fixation with EM fix solution (2.5% glutaraldehyde and 0.8% paraformaldehyde in PBS). Then the cells are sequentially incubated with 0.1 M imidazole for 2 h at 4℃, 1% osmium tetraoxide for 30 min at 4℃, PBS for 5 min, and 1% uranyl acetate overnight at 4℃. After staining, the cells are gradient dehydrated by ethanol and embedded in epoxy resin before being sectioned using Leica EM UC7 (~60 nm thick). Focus Ion Beam trimming and scanning EM images are acquired using a FEI Tecnai G2 20 Twin electron microscope and analyzed by Amria XImagePAQ Extension. The acquired EM images are reconstructed with Amria 3D and Chimera X.

**PAP activity assay**

PAP activity was determined in cell lysates by measuring the formation of DAG from PA (18:1) (Avanti® Polar lipids, Inc). Cells were harvested in 4°C by centrifugation at 800 rpm for 2 min, washed once with PBS and lysed by lysis buffer (50 mM Tris HCl pH 8.0, 10 mM β-mercaptoethanol, 5 μg/ml of aprotinin, leupeptin, and pepstatin, and 1x PhosSTOP phosphatase inhibitor cocktail). Then cell lysates were centrifuged at 13,000 rpm for 15 min in 4°C, the Lipins contained supernatant collected for the PAP reaction. In each reaction, 80 µg of total protein extract and 2 mM PA were added into reaction buffer (50 mM Tris HCl pH 8.0, 0.1 mM MgCl_2_, 5 mM EDTA, 0.1 mM ATP, 0.1 mM GTP) in a final volume of 100 µl. Reaction mixture were incubated for 16 h in 37°C. Lipids were extracted by methanal: dichloromethane = 1: 2 and separated the aqueous phase and organic phase by centrifuging at 13,000 rpm for 15 min in room temperature. Then organic-soluble lipid fraction was collected and dried by a rotary evaporator. The dry extracts were re-solubilized for DAG and PA measurement by LC-MS.

***In vivo* βCoV (MHV-A59)** **infection experiments**

Virus infection in mice is performed with wild type C57/BL6 male mice (4 weeks old) with intraperitoneal injection with MHV-A59 (p.f.u. = 2E3 for treatment experiments or 1E4 for combined treatment experiments and prophylactic experiments). The un-infected control group received a 200 μL volume of PBS via the same route. For drug delivery, the propranolol or isomers powders are dissolved in citrate buffer (pH = 3) and stored in -80 ℃ prior to diluted in 200 μL PBS for intraperitoneally delivery to mice. The Remdesivir stocks are dissolved in a mixed solution (DMSO, ethanol, PEG300 and PBS at 5:5:40:50) and stored in -20 ℃ prior to diluted in 200 μL PBS for intraperitoneal delivery. The control mice receive 200 μL PBS as placebos. The un-infected and βCoV infected mice are sacrificed at 7 days post infection. Other experiments are performed using littermates of 8-10 weeks old male mice. Mice are randomly separated into different groups for experiments.

**RNA extraction, RT-qPCR, and viral load determination**

Mouse tissues are mechanically disrupted by steel beads in 2 mL safe-lock tubes at 4 ℃. Tissue lysates are centrifuged at 12000 rpm for 5 min, prior to RNA extraction. Total RNA is extracted from tissue lysates and cell lysates by Trizol following manufacturer’s protocol. cDNA is generated using Superscript III RT kit (Transgen) and oligo-dT primers according to manufacturer’s protocol. Real-time PCR reactions are performed using the SYBR Green kit in the Archimed X4 fluorescence quantification system (Rocgene). *Scap^flox/flox^* mice used for *Tmem41b* transcripts determination are fasted from 6 p.m. on day one and refed from 6 p.m. on day two prior to sacrifice at 10 a.m. on day three. The primer sequences are listed in supplementary table 3.

**Histology**

Lung and liver samples are fixed in 4% PFA for 24 h and then transferred to 70% ethanol prior to dehydrating with graded ethanol and cleared with xylene solutions. Then tissues are embedded in paraffin, sectioned at 4 μm thickness and stained with Hematoxylin and Eosin (H/E) stain. Images are acquired by light microscopy in a double-blind manner and processed using ImageJ (Fuji).

**Molecular docking**

Molecular docking was utilized to elucidate the interaction between R-(+)-Propranolol and human Lipin1 catalytic domain. The crystal structure of R-(+)-Propranolol (ZINC code: 20240) was acquired from the ZINC database, with the 3D structure of human Lipin1 (UniProt code: Q14693) predicted by AlphaFold. Autodock software (version 4.2) was employed to perform the docking simulation, respectively. Conformations with the lowest docked energy were analyzed for the interactions between R-(+)-Propranolol and human Lipin1 catalytic domain. The 3D docking images were produced and processed by Pymol (version 2.0).

**Statistical analysis**

Quantification of IB and IF signals is performed by ImageJ (Fiji). Statistical analysis is performed by GraphPad Prism 8 and presented as Mean ± SEM of at least three independent experiments as indicated in the figure legends, and p-value is calculated with a two-tailed Student’s t test unless specifically noted. Results are considered significant when p < 0.05. Experiments of mice were performed in double blinded fashion with the random grouping.

**Antibodies**

| Rabbit anti-TMEM49/VMP1 (1: 1000) | Cell Signaling Technology | Cat# 12929 |
| --- | --- | --- |
| Rabbit anti-TMEM41B (1: 500) | Proteintech | Cat# 29270-1-AP |
| Rabbit anti-TMEM41A (1: 1000) | Proteintech | Cat# 20768-1-AP |
| Rabbit anti-LIPIN1 (1: 1000) | Cell Signaling Technology | Cat# 14906S |
| Rabbit anti-FASN (1: 1000) | Proteintech | Cat# 10624-2-AP |
| Rabbit anti-GM130 (1: 300) | Proteintech | Cat# 11308-1-AP |
| Rabbit anti-Tubulin (1: 5000) | Bioworld | Cat# AP0064 |
| Mouse anti-MHV N protein (1: 5000) | K. Holmes lab (U. of Colorado) | N/A |
| Mouse anti-dsRNA (1: 150) | Jena Bioscience | Cat# RNT-SCI-10010500 |
| Goat anti-Rabbit IgG (H+L) Secondary Antibody, HRP (1: 10000) | Thermo Fisher | Cat# 31460 |
| Goat anti-Mouse IgG (H+L) Secondary Antibody, HRP (1: 10000) | Pierce | Cat# 31430 |
| Alexa 488 Goat anti-Rabbit (1: 1000) | Thermo Fisher | Cat# A-11008 |
| Alexa 488 Goat anti-Mouse (1: 1000) | Thermo Fisher | Cat# A-11001 |
| Alexa 568 Goat anti-Rabbit (1: 1000) | Thermo Fisher | Cat# A-11011 |
| Alexa 568 Goat anti-Mouse (1: 1000) | Thermo Fisher | Cat# A-11031 |

**Chemicals, Critical Commercial Assay**

| Propranolol solution | Macklin | Cat# P902404 |
| --- | --- | --- |
| (±)-Propranolol hydrochloride | SIGMA | Cat# P0884 |
| S-(-)-Propranolol | Santa Cruz Biotechnology | Cat# sc-200153 |
| R-(+)-Propranolol | Perfemiker | Cat# PA80020 |
| Remdesivir | MedChemExpress | Cat# HY-104077 |
| Atenolol | Psaitong | Cat# A10837 |
| Acebutolol | MERDA | Cat# M020920 |
| PF-07321332 | Selleck | Cat# S9866 |
| Protease inhibitor | Roche | Cat# 4693132001 |
| TRIzol^TM^ Reagent | Thermo Fisher | Cat# 15596018 |
| KOD FX | Toyobo | Cat# KFX-101 |
| Alkyne choline | CONFLUORE | Cat# BCP-44 |
| Nile Red | Invitrogen | Cat# N1142 |
| TranScript One-Step gDNA Removal and cDNA Synthesis SuperMix | Transgen | Cat# AT311-03 |
| BCA Protein Assay Kit Pierce | Thermo Fisher | Cat# 23227 |
| EndoFree Maxi Plasmid Kit | TIANGEN | Cat# DP117 |
| OMEGA Gel Extraction Kit | OMEGA | D2500-02 |
| OMEGA Plasmid Mini Kit | OMEGA | D6943-02 |
| SuperReal PreMix Plus (SYBR Green) | TIANGEN | Cat# FP205 |
| Stbl3 Competent Cell | Tsingke | Cat# TSC06 |
| PA (18:1) | Avanti | Cat# 840875P |
| DAG (18:1) | Avanti | Cat# 800811C |

**Experimental Models: Cell lines, Organisms/Strains**

| HEK293T | ATCC | Cat# CRL-3216 |
| --- | --- | --- |
| Huh7 | JCRB Cell Ban | Cat# JCRB0403 |
| SpCas9 mice | Jackson Lab | Cat# JAX: 026556 |
| *Scap*^fl/fl^ mice | Jackson Lab | Cat# JAX: 004162 |

**Recombinant DNA**

| Plasmid: pX602 | Addgene | Cat# 61593 |
| --- | --- | --- |
| Plasmid: pX602-AAV-CRE-SgRNA | This paper | N/A |
| Plasmid: AAV-TBG-GFP | Addgene | Cat# 105535 |
| Plasmid: AAV-TBG-CRE | This paper | N/A |
| Plasmid: AAV-TBG-FLAG-TMEM41B | This paper | N/A |
| Plasmid: AAV-TBG-active SREBP1c | This paper | N/A |
| SARS-CoV-1 replicon | K. Holmes lab (U. of Colorado) | N/A |
| SARS-CoV-2 replicon | K. Holmes lab (U. of Colorado) | N/A |
| Sanger Arrayed Whole Genome Lentiviral CRISPR Library Human | Sigma-Aldrich | Cat# HSANGERG |

**Software and Algorithms**

| ImageJ (Fiji) | NIH, Schindelin et al, 2012 | RRID: SCR_0030. 70 |
| --- | --- | --- |
| GraphPad Prism8 | GraphPad Software | N/A |
| Benchling | Benchling | https://benchling.com/ |
| Pymol | Schrödinger, LLC | https://pymol.org |

**Sequences of oligonucleotides for genotyping.**

| Primer sequences for genotyping | | |
| --- | --- | --- |
|  | Forward (5’ - 3’) | Reverse (5’ - 3’) |
| spCas9 KI | AGGAAGCACTTGCTCTCCCA | CTGTTCAATTCCCCTGCAGGAC  GAAACTTTCGGAGCGCGCC |
| *Scap fl/fl* | GCTCTGCGCATCCTATCCAATTCC | CAGCCGGCAAGTAACAAGGGATCC |

**Sequences of oligonucleotides for sgRNA constructs.**

| Sequences of oligonucleotides for sgRNA constructs | |
| --- | --- |
|  | Guide sequence (5’ - 3’) |
| LacZ-sg | TGCGAATACGCCCACGCGAT |
| Mouse *Tmem41b*-sg | GGAACCCGGAGAGTATACTG |
| Mouse *Lipin1-*sg | GGAAGAAGAGAAGGAAAAGG |
| Human *TMEM41B-*sg | GTCGCCGAACGATCGCAGTT |

**Sequences of oligonucleotides for qPCR.**

| Sequences of oligonucleotides for qPCR | | |
| --- | --- | --- |
| Primers for mouse sample | | |
|  | Forward (5’ - 3’) | Reverse (5’ - 3’) |
| *β-Actin* | GGCTGTATTCCCCTCCATCG | CCAGTTGGTAACAATGCCATGT |
| *Fasn* | TCCAAGACTGACTCGGCTACTGAC | GCAGCCAGGTTCGGAATGCTA TC |
| *Tmem41b* | GATATGGATGACGCCAAGGCT | CAAACAAACAAGGAATAAGGCAAG |
| *Tmem41a* | ATACCTGCTCTCCACGCGGC | GCAGCTCTGCCAGATCGGAG |
| *Vmp1* | AGAAGAAGAGAAGGGATCGG | TTGGTGCGCTCCTTCCACAT |
| *Tnfa* | GGTGCCTATGTCTCAGCCTCTT | CGATCACCCCGAAGTTCAGTA |
| *Il6* | CACAGAGGATACCACATCCCAACA | TCCACGATTTCCCAGAGAACA |
| MHV-N | CAGATCCTTGATGATGGCGTAGT | AGAGTGTCCTATCCCGACTTTCTC |
